# Genome-wide screening reveals producer-cell modifications that improve virus-like particle production and delivery potency

**DOI:** 10.1101/2025.10.07.681010

**Authors:** Diana Ly, Hyewon Jang, Adhiraj Goel, Arnav Singh, Aditya Raguram

## Abstract

Engineered virus-like particles (eVLPs) are promising vehicles for transient delivery of gene editing agents. While extensive particle engineering has yielded efficient eVLPs, it remains underexplored whether engineering the cells used to produce eVLPs could further improve eVLP properties. We developed a genome-wide screening approach to systematically investigate how genetic perturbations in producer cells influence eVLP production. This approach generates eVLPs loaded with guide RNAs that identify the genetic perturbation in the cell that produced a particular particle; the abundance of each guide RNA in eVLPs therefore reflects how the corresponding genetic perturbation influences eVLP production or cargo loading. We applied this approach to identify several genes that regulate eVLP cargo expression and loading into particles during the production process. Leveraging these insights, we engineered producer cells that support increased eVLP cargo packaging and a 2- to 9-fold increase in eVLP delivery potency across several cargo, particle, and target-cell types in cultured cells and in mice. Our findings suggest the potential of producer-cell engineering as a useful strategy for improving the utility of eVLPs and related delivery methods.

## Introduction

Methods for safely and efficiently delivering macromolecules into cells in culture (*in vitro*) and within the body (*in vivo*) are required for many therapeutic strategies, including gene editing therapies^1–5^. Recently, virus-like particles (VLPs) have emerged as promising vehicles for therapeutic macromolecule delivery that combine key advantages of canonical viral and non-viral delivery strategies^1,6^. In typical VLPs, viral scaffold proteins are used to package and deliver cargo proteins, ribonucleoproteins (RNPs), or mRNAs instead of viral genetic material, which leads to transient cargo delivery into recipient cells^1,6^. Several VLP-based delivery methods have proven particularly effective for gene editing applications^7–23^ and offer an ideal combination of efficient on-target editing, minimal off-target editing, and programmable cell-type targeting capabilities. Current state-of-the-art VLPs, including engineered VLPs (eVLPs)^7,8,22^, have achieved efficient *in vitro* gene editing in primary human cells and therapeutic *in vivo* gene editing in the mouse liver and retina, highlighting the utility of eVLP delivery for research and potential therapeutic applications.

While extensive engineering of eVLP components and particle architectures has afforded efficient eVLPs^7,8,22^, further improvements to eVLP properties are still required to maximize their utility. In particular, improving eVLP cargo packaging per particle or overall particle yield from the eVLP production process would enable more efficient gene editing with lower eVLP doses and simplify the production of eVLPs for large-scale studies. Standard methods for producing eVLPs involve the use of human producer cells to express eVLP components and assemble functional particles^7,8,22^, suggesting that producer cells likely play an important role in the eVLP production process. Indeed, producer-cell engineering has previously been used to improve the production of lentiviral vectors and adeno-associated viral vectors^24–29^. However, previous studies have focused solely on particle engineering to improve eVLP properties, and none have explored whether producer-cell engineering could improve eVLP properties beyond what can be achieved through particle engineering alone.

In this study, we systematically investigated how genetic perturbations in producer cells influence eVLP production and applied our findings to improve eVLP production and delivery potency. Using an unbiased genome-wide screening approach in which eVLP-packaged guide-RNA cargos directly correspond to producer-cell genetic perturbations, we identified several genes that impact eVLP production, including genes that are uniquely relevant for packaging RNP cargos into genome-free eVLPs. Guided by these observations, we engineered producer cells that support improved eVLP production and delivery potency across several cargo, particle, and target-cell types in cultured cells and in mice. Our results lay a foundation for using producer-cell engineering to improve the utility of eVLPs and related bioparticle-based delivery methods.

## Results

### Genome-wide screening reveals producer-cell perturbations that influence eVLP production

We began by developing an unbiased genome-wide screening approach to identify genetic perturbations in producer cells that influence eVLP production (Fig. 1a). We envisioned a general scheme in which eVLP production is initiated from a pool of genetically perturbed producer cells. Each producer cell in this pool expresses a Cas9-based genome modifying agent and a single copy of a guide RNA (sgRNA) that directs the installation of a specific genetic perturbation into that particular cell (Fig. 1a). Because eVLPs package Cas9-based cargos, the perturbation-inducing sgRNA expressed in a given producer cell is also loaded into any particles produced by that particular cell (Fig. 1a). Therefore, the abundance of a particular sgRNA in eVLPs reflects how the corresponding genetic perturbation influences eVLP production or cargo loading: sgRNAs that are more abundant identify perturbations that increase eVLP production or cargo loading, while sgRNAs that are less abundant identify perturbations that decrease eVLP production or cargo loading (Fig. 1a).

**Fig. 1.**
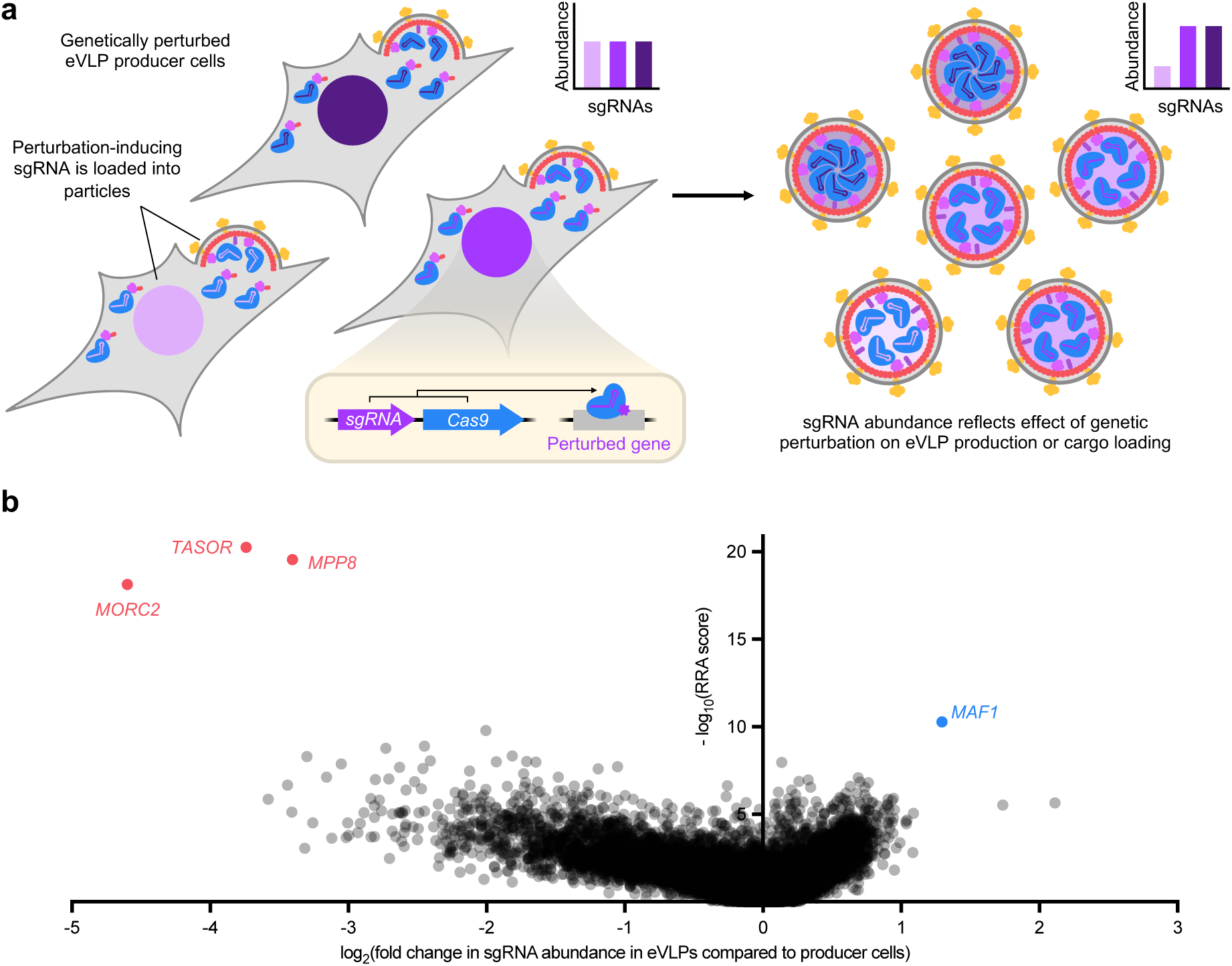
Genome-wide screening reveals genetic perturbations that influence eVLP production. **a**, Overview of the producer-cell screening approach. eVLP production is initiated from a pool of genetically perturbed producer cells, and each eVLP packages sgRNAs that identify the specific genetic perturbation in the cell that produced that particular eVLP. The abundance of a particular sgRNA in eVLPs compared to producer cells reflects how the corresponding genetic perturbation influences eVLP production or cargo loading. **b**, Gene-level phenotypes from the genome-wide knockout screen for cargo-loaded eVLP production. Each gene is represented as a single dot whose x-coordinate reflects the median fold change in sgRNA abundance in eVLPs compared to producer cells for all sgRNAs targeting that particular gene, and whose y-coordinate reflects the MAGeCK RRA score associated with that particular gene (see Methods). Data reflect n=2 screen replicates. See also Extended Data Fig. 1c and Supplementary Table 6.

We used this general approach to perform a genome-wide knockout screen for cargo-loaded eVLP production in Gesicle 293T producer cells, a standard cell line used for eVLP production^7,8,22^. We transduced producer cells at a low multiplicity of infection (MOI) with a lentiviral sgRNA library containing 98,077 unique sgRNAs targeting 19,707 genes across the human genome^30^ (Extended Data Fig. 1a). Each vector in the library also encoded constitutive expression of Cas9 nuclease. After selecting for Cas9 expression and expanding the transduced cells to generate a pool of single-knockout producer cells, we initiated fourth-generation (v4) adenine base editor (ABE)-eVLP production from these cells (Extended Data Fig. 1a). Importantly, to initiate eVLP production, we only transfected plasmids encoding the expression of v4 ABE-eVLP protein components: (1) the gag–ABE (capsid–cargo) fusion, (2) the Moloney murine leukemia virus (MMLV) gag–pro–pol polyprotein, and (3) the vesicular stomatitis virus G (VSV-G) envelope protein. No additional sgRNA-encoding plasmid was transfected, ensuring that all eVLPs produced from a given producer cell packaged cargo containing the specific knockout-inducing sgRNA that was already introduced into that producer cell by lentiviral transduction (Fig. 1a and Extended Data Fig. 1a).

Following eVLP production, we isolated RNA from purified eVLPs and compared this eVLP-packaged RNA to cellular RNA that was isolated from the single-knockout producer-cell pool immediately prior to eVLP production (Fig. 1a and Extended Data Fig. 1a). To compare sgRNA abundances across both populations, we subjected both RNA pools to template-switching reverse transcription^31^, which enabled amplification and sequencing of the spacer sequence at the 5’ end of each sgRNA molecule (Extended Data Fig. 1b). sgRNAs that are more abundant in the eVLP-packaged RNA compared to the producer-cell RNA identify gene knockouts that increase eVLP production or cargo loading, while sgRNAs that are less abundant identify gene knockouts that decrease eVLP production or cargo loading (Fig. 1a and Extended Data Fig. 1a).

By aggregating the results for all sgRNAs targeting the same gene into gene-level phenotypes, we determined the effect of each producer-cell gene knockout on eVLP production or cargo loading (Fig. 1b and Extended Data Fig. 1c). We observed that the majority of gene knockouts did not substantially impact eVLP production or cargo loading, some knockouts decreased eVLP production or cargo loading, and very few knockouts increased eVLP production or cargo loading (Fig. 1b). These results are consistent with a model in which there are more genes in producer cells that positively regulate eVLP production than genes that negatively regulate eVLP production. Taken together, the results of this genome-wide knockout screen illuminate the producer-cell determinants of eVLP production and nominate several gene knockouts (“hits”) that appear to strongly impact eVLP production or cargo loading (Fig. 1b).

### Knockout of HUSH-associated genes negatively impacts eVLP sgRNA packaging in the presence of lentiviral Cas9 expression

Next, we sought to characterize the gene knockouts that most strongly decreased eVLP production or cargo loading. While these negative hits might not be immediately applicable for improving eVLP manufacturing, we reasoned that characterizing these gene knockouts would reveal important insights into how producer-cell factors influence eVLPs, which could inform alternative strategies for improving eVLP production. The three strongest negative hits from our screen—*MPP8*, *TASOR*, and *MORC2*—are all associated with the human silencing hub (HUSH) complex, which is responsible for epigenetically silencing intronless transgenes, including retroviral elements^32,33^. We first individually validated each negative hit by measuring the relative sgRNA abundance in eVLPs produced from *MPP8*-, *TASOR*-, or *MORC2*-knockout cells (hereafter collectively referred to as HUSH-knockout cells) compared to eVLPs produced from standard producer cells. To independently generate each knockout cell line, we mimicked the screen conditions and transduced Gesicle cells with a lentiviral vector encoding constitutive expression of Cas9 nuclease and an appropriate targeting sgRNA, selected cells for Cas9 expression, and isolated a polyclonal population of Cas9-edited cells (Extended Data Fig. 2a). To control for the effects of Cas9 expression and Cas9 nuclease-mediated indel generation on producer cells, we also applied the same procedure to generate modified Gesicle cells using a sgRNA targeting *AAVS1*, a human safe harbor locus^34^. After knockout generation, we initiated v4 ABE-eVLP production from each cell line by transfecting plasmids encoding the expression of all required components, including a sgRNA targeting the human *BCL11A* +58 enhancer locus^35^ (Extended Data Fig. 2a). We detected by RT-qPCR a 1.5- to 3.8-fold decrease in sgRNA abundance in ABE-eVLPs produced from HUSH-knockout cells compared to ABE-eVLPs produced from *AAVS1*-knockout cells (Fig. 2a), confirming the results of our screen. Additionally, when we transduced HEK293T cells with a range of doses of purified ABE-eVLPs and measured adenine base editing efficiencies at the *BCL11A* +58 enhancer target locus via high-throughput sequencing (Extended Data Fig. 2a), we observed that ABE-eVLPs produced from HUSH-knockout cells exhibited a 2.3- to 4.4-fold decrease in *in vitro* delivery potency (EC_50_) compared to ABE-eVLPs produced from *AAVS1*-knockout cells (Fig. 2b). These results indicate that knockout of HUSH-associated genes *MPP8*, *TASOR*, or *MORC2* in producer cells negatively impacts eVLP production or cargo loading.

**Fig. 2.**
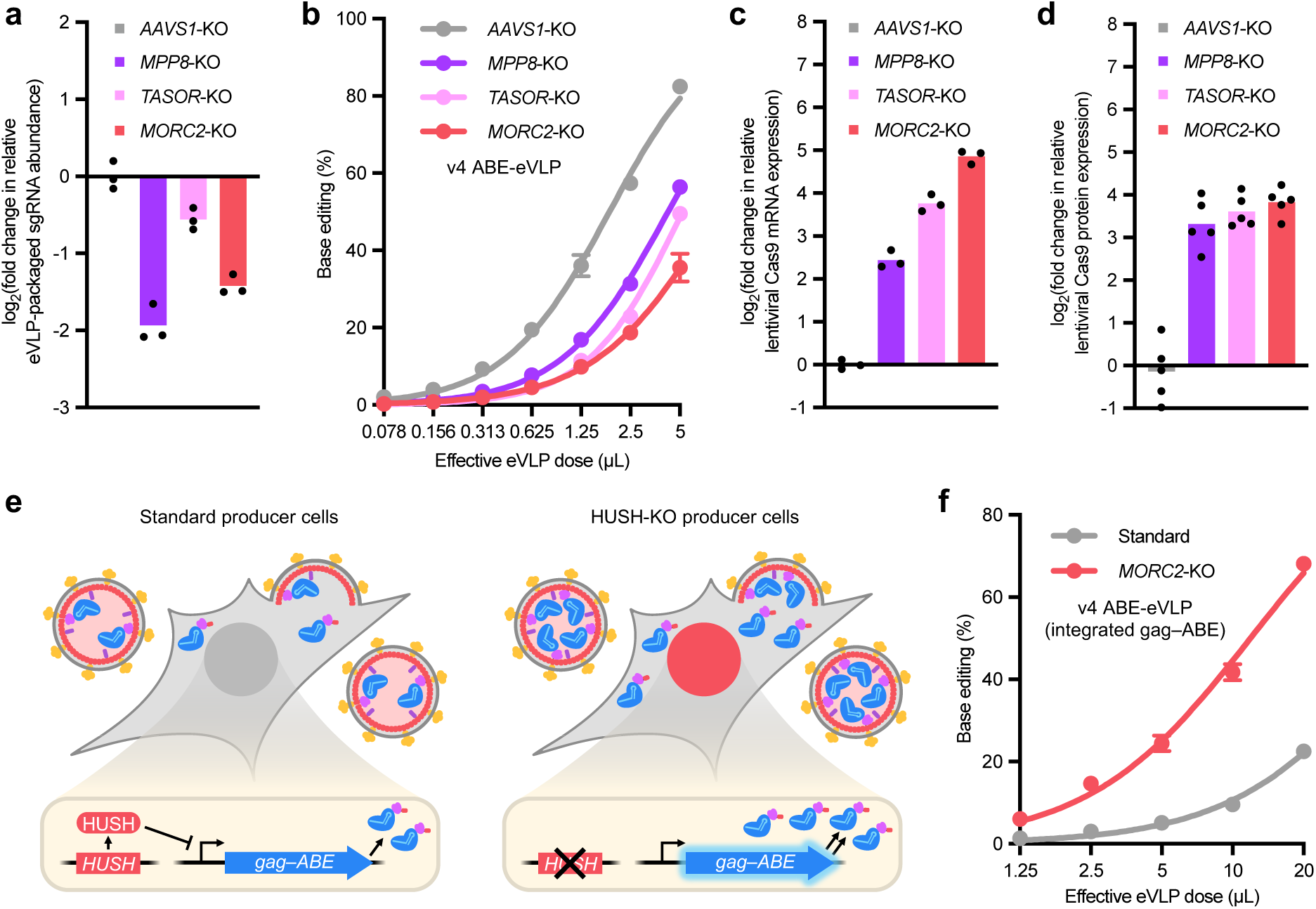
Knockout of HUSH-associated genes negatively impacts eVLP sgRNA packaging and delivery potency. **a**, Fold change in eVLP-packaged sgRNA abundance measured by RT-qPCR, normalized relative to sgRNA abundance in eVLPs produced from *AAVS1*-knockout cells. Bars reflect the mean of n=3 replicates, and dots represent individual replicate values. **b**, Comparison of v4 ABE-eVLPs produced from *AAVS1*-knockout or HUSH-knockout cells across a range of eVLP doses. Adenine base editing efficiencies at position A_7_ of the *BCL11A* +58 enhancer site in HEK293T cells are shown. **c**, Fold change in producer-cell Cas9 mRNA abundance measured by RT-qPCR, normalized relative to Cas9 mRNA abundance in *AAVS1*-knockout cells. Bars reflect the mean of n=3 replicates, and dots represent individual replicate values. **d**, Fold change in producer-cell Cas9 protein abundance measured by Western blot, normalized relative to Cas9 protein abundance in *AAVS1*-knockout cells. Bars reflect the mean of n=5 replicates, and dots represent individual replicate values. See also Extended Data Fig. 3a. **e**, Schematic of standard producer cells versus HUSH-knockout producer cells, both containing genomically integrated gag–ABE expression cassettes. In standard cells, epigenetic silencing by the HUSH complex limits gag–ABE expression. In HUSH-knockout cells, lack of HUSH-mediated silencing leads to increased gag–ABE expression and loading into eVLPs. **f**, Comparison of v4 ABE-eVLPs produced from standard or *MORC2*-knockout Gesicle cells, both containing genomically integrated gag–ABE expression cassettes. Adenine base editing efficiencies at position A_5_ of the *HEK2* site in HEK293T cells are shown. In **b** and **f**, dots and error bars represent mean ± s.d. of n=3 biological replicates. Data were fit to four-parameter logistic curves using nonlinear regression.

We hypothesized that these HUSH-associated genes might impact eVLP production or cargo loading by modulating the expression of the lentivirally integrated Cas9 nuclease that was used to generate the gene knockouts. To test this hypothesis, we measured the relative abundance of Cas9 mRNA or protein in each knockout cell line by RT-qPCR or Western blot respectively. We observed a 5.4- to 29-fold increase in cellular Cas9 mRNA expression and a 10- to 14-fold increase in Cas9 protein expression in HUSH-knockout cells compared to *AAVS1*-knockout cells (Fig. 2c, d and Extended Data Fig. 3a). These results suggest a model in which HUSH-associated gene products natively silence lentivirally integrated Cas9 expression in *AAVS1*-knockout producer cells, but in HUSH-knockout cells, Cas9 is no longer efficiently silenced (Extended Data Fig. 3b). Therefore, during eVLP production, excess Cas9 protein in HUSH-knockout producer cells likely competes for sgRNA binding and prevents sgRNA molecules from efficiently binding to gag–ABE, which ultimately decreases both sgRNA packaging into eVLPs and eVLP delivery potency (Extended Data Fig. 3b). To further investigate this phenomenon, we used v4 Cas9-eVLP transduction instead of Cas9/sgRNA lentiviral transduction to generate HUSH-knockout Gesicle cells that lacked any integrated Cas9 transgene (Extended Data Fig. 2b). Notably, we did not detect any change in sgRNA abundance by RT-qPCR in ABE-eVLPs produced from these Cas9-free HUSH-knockout cells compared to standard producer cells (Extended Data Fig. 4a). These results suggest that, consistent with our model above, both lentivirally integrated Cas9 expression and knockout of HUSH-associated genes are required to decrease sgRNA abundance in eVLPs. Collectively, these data and our proposed model rationalize the negative performance of HUSH-associated gene knockouts in our screen.

### Knockout of HUSH-associated genes increases eVLP protein cargo expression

Interestingly, we noticed that ABE-eVLPs produced from Cas9-free HUSH-knockout cells still exhibited a 2.6- to 3.9-fold decrease in delivery potency compared to ABE-eVLPs produced from standard Gesicle cells (Extended Data Fig. 4b), suggesting that some eVLP property other than sgRNA abundance was negatively impacted due to Cas9-free HUSH-associated gene knockout. We hypothesized that these HUSH-associated genes might also modulate the expression of eVLP components from plasmids transfected during the eVLP production process. To investigate this possibility, we performed Western blots to quantify eVLP cargo (gag–ABE) expression in producer cells following transfection of eVLP production plasmids, which revealed a 2.1- to 4.4-fold increase in gag–ABE protein expression in HUSH-knockout cells compared to Gesicle cells (Extended Data Fig. 4c, d). We reasoned that this increase in eVLP cargo expression might disrupt the optimal eVLP component stoichiometry (gag–ABE:gag–pro–pol ratio), which was previously found to be an important determinant of eVLP potency^8^. Consistent with this model, we observed that reducing the transfected gag–ABE:gag–pro–pol plasmid ratio from 25:75 to 10:90 or 5:95 restored the potency of ABE-eVLPs produced from *MORC2*-knockout cells back to standard levels (Extended Data Fig. 4e). These results indicate that the increase in eVLP cargo expression in HUSH-knockout producer cells and corresponding disruption of the optimal eVLP component stoichiometry can be counteracted simply by reducing the amount of transfected eVLP cargo plasmid, thereby effectively restoring the optimal stoichiometry.

Given our finding that knockout of HUSH-associated genes increases eVLP cargo expression, we wondered whether HUSH-knockout producer cells could in fact improve eVLP production in specific contexts. In particular, certain applications of eVLPs, including our recently described barcoded eVLP evolution system^22^, require expressing gag–ABE from genomically integrated cassettes rather than transfected plasmids, which results in reduced gag–ABE protein levels and suboptimal component stoichiometry. We reasoned that the use of HUSH-knockout producer cells should increase gag–ABE expression from genomically integrated cassettes, which in this case would improve rather than disrupt the eVLP component stoichiometry (Fig. 2e). Consistent with this model, we observed that ABE-eVLPs produced from *MORC2*-knockout cells with integrated gag–ABE expression exhibited a 4.6-fold increase in delivery potency compared to ABE-eVLPs produced from standard cells with integrated gag–ABE expression (Fig. 2f). These results indicate that HUSH-knockout producer cells support improved eVLP delivery potency in situations in which gag–cargo expression is limiting. Altogether, our results reveal a previously unknown role for HUSH-associated genes in modulating eVLP cargo expression in producer cells and highlight the importance of optimizing eVLP cargo expression and component stoichiometry to maximize eVLP potency.

### MAF1-knockout producer cells improve sgRNA packaging into eVLPs

Next, we sought to characterize the gene knockouts from our screen that most strongly increased eVLP production or cargo loading. The single strongest positive hit from our screen was *MAF1*, a repressor of RNA Pol III transcription^36–38^. Because all sgRNAs in our experiments were expressed from an RNA Pol III-transcribed U6 promoter, we reasoned that knockout of *MAF1* in producer cells likely leads to increased sgRNA transcription and loading into eVLP-packaged cargos (Fig. 3a).

**Fig. 3.**
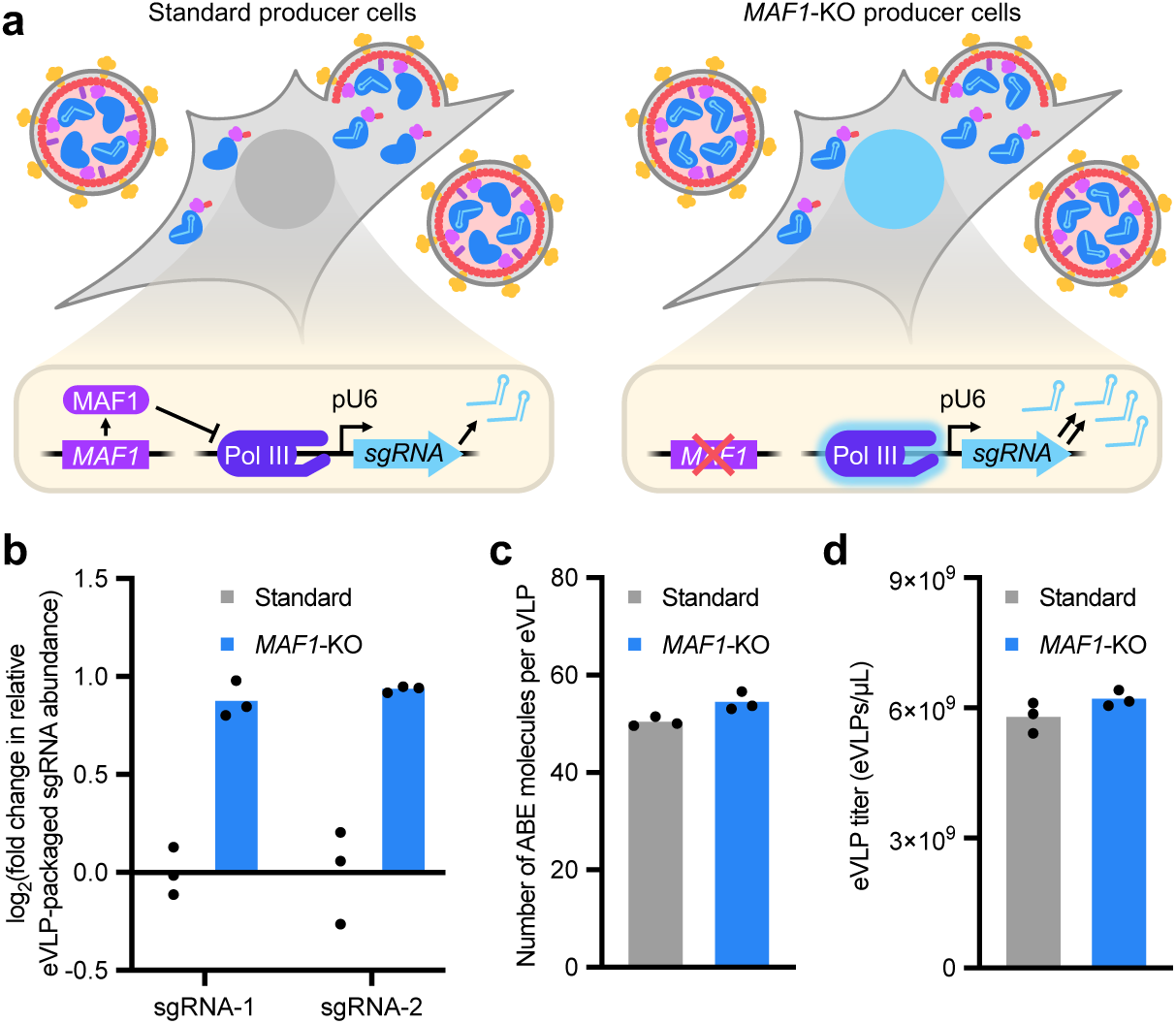
*MAF1*-knockout producer cells improve eVLP sgRNA packaging. **a**, Schematic of standard producer cells versus *MAF1*-knockout producer cells. In standard cells, MAF1 represses RNA Pol III transcription, which limits producer-cell sgRNA expression. In *MAF1*-knockout producer cells, RNA Pol III is no longer subjected to MAF1-mediated repression, which increases sgRNA expression and loading into eVLP-packaged RNPs. **b**, Fold change in eVLP-packaged sgRNA abundance measured by RT-qPCR, normalized relative to sgRNA abundance in eVLPs produced from standard Gesicle cells. sgRNA-1 and sgRNA-2 denote two different packaged sgRNA sequences. **c–d**, Quantification of ABE molecules per eVLP (**c**) and eVLP particle titer (**d**) by anti-Cas9 and anti-MLV p30 ELISA (see Methods). In **b–d**, bars reflect the mean of n=3 replicates, and dots represent individual replicate values.

To test this hypothesis, after generating *MAF1*-knockout Gesicle cells using v4 Cas9-eVLPs (Extended Data Fig. 2b), we initiated v4 ABE-eVLP production from these cells by transfecting plasmids encoding the expression of all required components. We detected by RT-qPCR a 2-fold increase in sgRNA abundance in ABE-eVLPs produced from *MAF1*-knockout cells compared to ABE-eVLPs produced from standard Gesicle cells (Fig. 3b). Additionally, we observed that ABE-eVLPs produced from *MAF1*-knockout cells were produced at similar particle titers and packaged similar numbers of ABE cargo proteins per particle compared to ABE-eVLPs produced from standard Gesicle cells (Fig. 3c, d), suggesting that the use of *MAF1*-knockout cells increases sgRNA packaging alone without affecting protein packaging or particle yield. Collectively, these data confirm our knockout screen results and indicate that *MAF1*-knockout producer cells support increased sgRNA loading into eVLPs.

### MAF1-knockout producer cells support improved eVLP delivery potency in cell culture and in mice

Because increased sgRNA loading leads to an increased number of active RNP cargo molecules packaged into eVLPs (Fig. 3a), we reasoned that ABE-eVLPs produced from *MAF1*-knockout cells should exhibit increased delivery potency compared to ABE-eVLPs produced from standard Gesicle cells. Encouragingly, we observed that v4 ABE-eVLPs produced from *MAF1*-knockout cells exhibited a 2.5-fold increase in delivery potency (EC_50_) at the *BCL11A* +58 enhancer locus in HEK293T cells compared to v4 ABE-eVLPs produced from standard Gesicle cells (Fig. 4a). v4 ABE-eVLPs produced from *MAF1*-knockout cells also exhibited a 2.2- and 2.1-fold increase in delivery potency at the *GAPDH* and *HIRA* genomic loci in HEK293T cells, respectively (Fig. 4b, c), indicating that the eVLP potency improvement conferred by the use of *MAF1*-knockout producer cells is not restricted to any particular sgRNA sequence or target genomic locus. Additionally, we assessed whether the use of *MAF1*-knockout producer cells could also improve the delivery potency of the more recently developed v5 ABE-eVLPs, which use laboratory evolved capsid proteins that were optimized to improve BE RNP cargo packaging^22^. We observed that v5 ABE-eVLPs produced from *MAF1*-knockout cells exhibited a 2.2- to 9.1-fold increase in delivery potency compared to v5 ABE-eVLPs produced from standard cells across six distinct target genomic loci in mouse Neuro-2a cells (Fig. 4d–i). Together, these results indicate that the increased sgRNA packaging in ABE-eVLPs produced from *MAF1*-knockout cells leads to corresponding improvements in eVLP delivery potency *in vitro*.

**Fig. 4.**
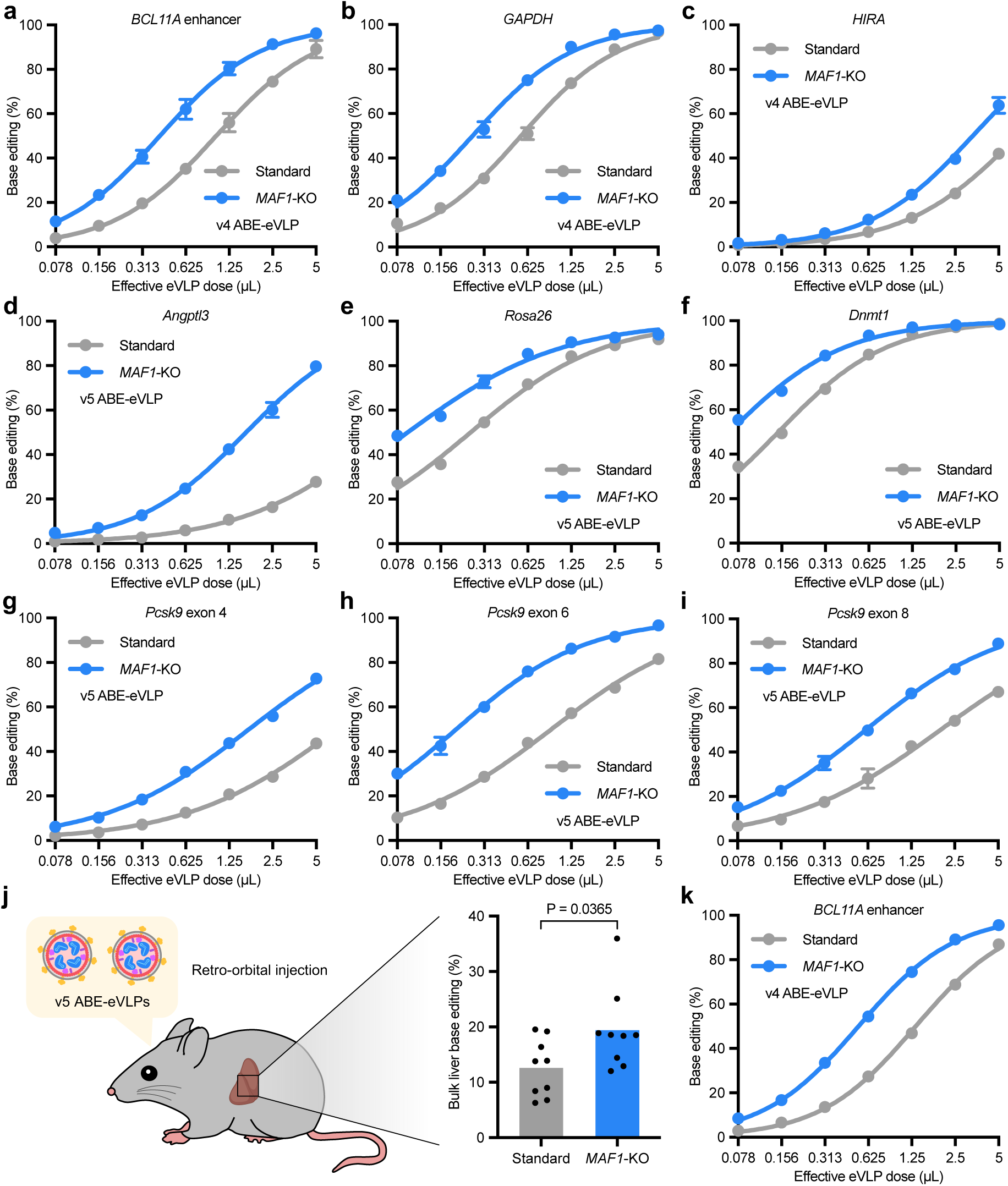
*MAF1*-knockout producer cells improve eVLP delivery potency in cultured cells and in mice. **a–c**, Comparison of v4 ABE-eVLPs produced from standard or *MAF1*-knockout Gesicle cells across a range of eVLP doses in HEK293T cells. Adenine base editing efficiencies are shown at position A_7_ of the *BCL11A* +58 enhancer site (**a**), position A_7_ of the *GAPDH* site (**b**), and position A_5_ of the *HIRA* site (**c**). **d–i**, Comparison of v5 ABE-eVLPs produced from standard or *MAF1*-knockout Gesicle cells across a range of eVLP doses in mouse Neuro-2a cells. Adenine base editing efficiencies are shown at position A_8_ of the *Angptl3* exon 7 splice acceptor site (**d**), position A_4_ of the *Rosa26* site, position A_9_ of the *Dnmt1* site (**f**), position A_6_ of the *Pcsk9* exon 4 splice acceptor site (**g**), position A_4_ of the *Pcsk9* exon 6 splice donor site (**h**), and position A_8_ of the *Pcsk9* exon 8 splice acceptor site (**i**). **j**, Schematic of mouse experiments and adenine base editing efficiencies at position A_6_ of the *Pcsk9* exon 1 splice donor site in bulk mouse liver tissue. Each mouse was injected with 15 µL of 1500-fold concentrated purified v5 ABE-eVLPs containing 3×10^11^ eVLPs as determined by anti-MLV p30 ELISA (see Methods). Bars reflect the mean of n=9 mice per condition, and dots represent individual mouse values. P-value was calculated using a two-sided unpaired t test. **k**, Comparison of v4 ABE-eVLPs produced from standard or *MAF1*-knockout HEK293T/17 cells across a range of eVLP doses in HEK293T cells. Adenine base editing efficiencies at position A_7_ of the *BCL11A* +58 enhancer site are shown. In **a–i** and **k**, dots and error bars represent mean ± s.d. of n=4 (**a**) or n=3 (**b–i**, **k**) biological replicates. Data were fit to four-parameter logistic curves using nonlinear regression.

To further explore the utility of *MAF1*-knockout producer cells, we investigated whether their use could also improve *in vivo* eVLP delivery efficiencies. We produced v5 ABE-eVLPs targeting the *Pcsk9* exon 1 splice donor^8,39^ using either standard or *MAF1*-knockout Gesicle cells, and we injected these ABE-eVLPs retro-orbitally into adult mice at a low, sub-saturating eVLP dose (Fig. 4j). We observed that v5 ABE-eVLPs produced from *MAF1*-knockout cells achieved 19.4% average *in vivo* base editing at the *Pcsk9* locus in bulk liver tissue, while v5 ABE-eVLPs produced from standard cells administered at the same dose only achieved 12.6% bulk liver editing (Fig. 4j, *P* = 0.0365, two-sided unpaired *t* test). Collectively, these data indicate that *MAF1*-knockout producer cells support improved eVLP potency across different target genomic loci and target-cell types both *in vitro* and *in vivo*, demonstrating the benefits of using *MAF1*-knockout cells instead of standard cells for eVLP production.

### MAF1-knockout producer cells support improved delivery potency across various cargo, particle, and producer-cell types

We next sought to further evaluate the ability of *MAF1*-knockout producer cells to support improved eVLP production and potency in other various contexts. First, we investigated whether knockout of *MAF1* could improve eVLP production from HEK293T/17 cells, another cell line that is commonly used for viral vector production^8,22,40^. Following the generation of *MAF1*-knockout HEK293T/17 cells using v4 Cas9-eVLPs, we observed that v4 ABE-eVLPs produced from *MAF1*-knockout HEK293T/17 cells exhibited a 2.5-fold increase in *in vitro* delivery potency compared to those produced from standard HEK293T/17 cells (Fig. 4k). These results indicate that the improvements conferred by *MAF1* knockout are generalizable beyond the specific cell line (Gesicle 293T) in which our knockout screen was conducted. Additionally, our findings suggest that knockout of *MAF1* might similarly benefit eVLP production from HEK293T/17 clones that have been adapted for growth in suspension culture, which are particularly well suited for large-scale bioparticle manufacturing^41^.

Second, we assessed whether *MAF1*-knockout producer cells could improve the potency of eVLPs that package different Cas9-based RNP cargos. We observed that v4 eVLPs packaging either Cas9 nuclease, ABE-SpCas9-NG^42^, or TadCBEa^43^ cargos exhibited a 2.0- to 2.5-fold improvement in delivery potency when produced from *MAF1*-knockout cells compared to Gesicle cells (Fig. 5a–e).

**Fig. 5.**
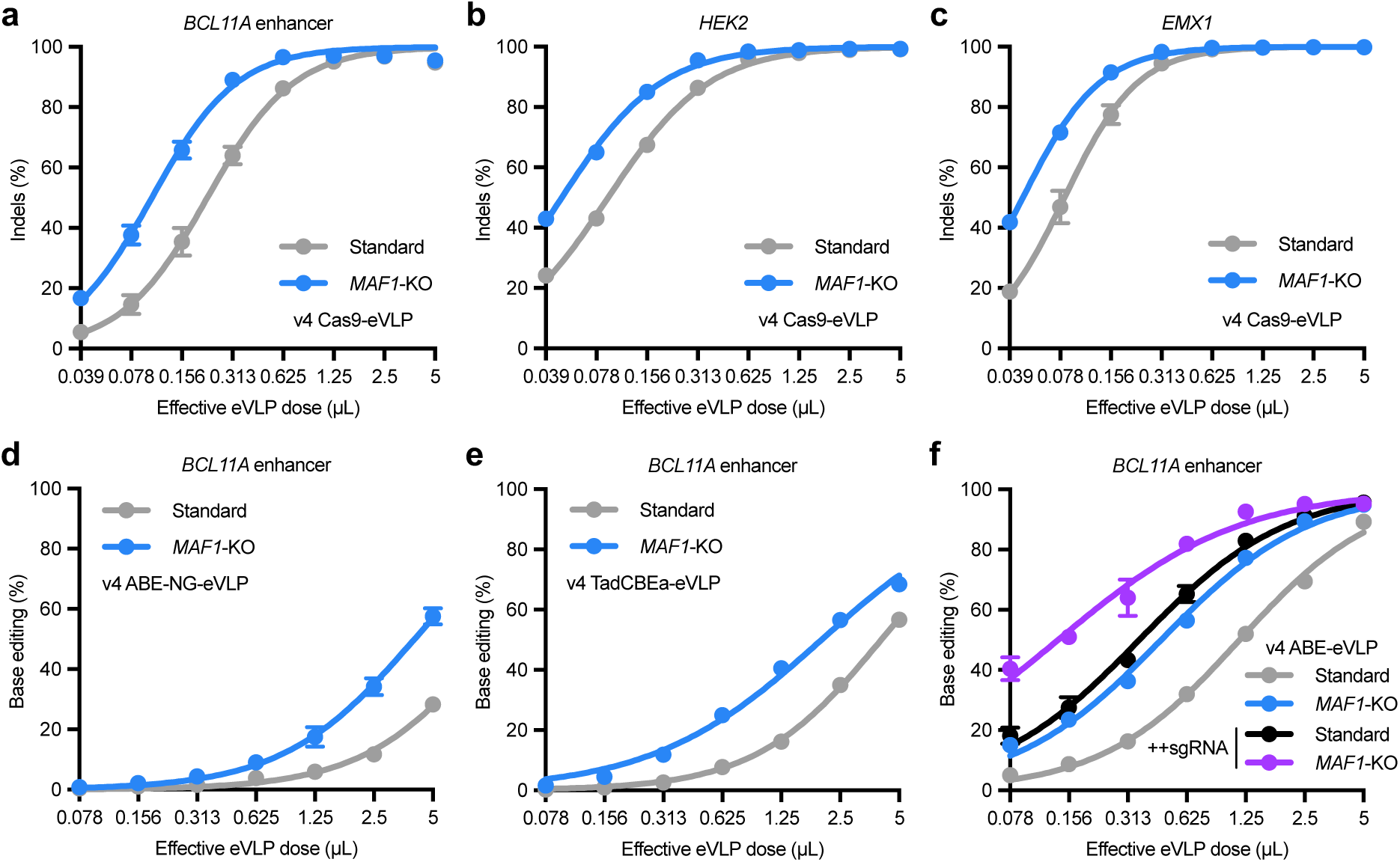
*MAF1*-knockout producer cells improve eVLP delivery potency across cargo types. **a–c**, Comparison of v4 Cas9-eVLPs produced from standard or *MAF1*-knockout Gesicle cells across a range of eVLP doses in HEK293T cells. Indel frequencies are shown at the *BCL11A* +58 enhancer site (**a**), *HEK2* site (**b**), and *EMX1* site (**c**). **d**, Comparison of v4 ABE-NG-eVLPs produced from standard or *MAF1*-knockout Gesicle cells across a range of eVLP doses. Adenine base editing efficiencies at position A_7_ of the *BCL11A* +58 enhancer site in HEK293T cells are shown. **e**, Comparison of v4 TadCBEa-eVLPs produced from standard or *MAF1*-knockout Gesicle cells across a range of eVLP doses. Cytosine base editing efficiencies at position C_6_ of the *BCL11A* +58 enhancer site in HEK293T cells are shown. **f**, Comparison of v4 ABE-eVLPs or v4 ABE-eVLPs with additional sgRNA-expressing plasmids (++sgRNA, see Extended Data Fig. 7) produced from standard or *MAF1*-knockout Gesicle cells across a range of eVLP doses. Adenine base editing efficiencies at position A_7_ of the *BCL11A* +58 enhancer site in HEK293T cells are shown. **a–f**, Dots and error bars represent mean ± s.d. of n=3 biological replicates. Data were fit to four-parameter logistic curves using nonlinear regression.

Third, we explored whether *MAF1*-knockout producer cells could also improve the performance of non-eVLP delivery vehicles, including several recently reported RNP-packaging VLP- and vesicle-based systems. We evaluated four different non-eVLP systems that use various particle scaffolds and cargo loading mechanisms (Extended Data Fig. 5a): (1) ENVLPE^+^ (engineered nucleocytosolic vehicles for loading of programmable editors)^10^, (2) miniEDV (minimal enveloped delivery vehicles)^44^, (3) RIDE VLPs^16^, and (4) VEDIC (VSV-G plus EV-Sorting Domain-Intein-Cargo)^45^ particles. We observed that ABE-packaging ENVLPE^+^ particles produced from *MAF1*-knockout cells exhibited a 2.2-fold increase in delivery potency compared to the same particles produced from standard Gesicle cells, which was nearly identical to the improvement we observed for v4 ABE-eVLPs (Extended Data Fig. 5b, c). Prime editor (PE)-packaging ENVLPE^+^ particles produced from *MAF1*-knockout cells also exhibited a 2.1-fold increase in delivery potency compared to the same particles produced from standard Gesicle cells (Extended Data Fig. 5d). Additionally, we observed that Cas9-packaging miniEDVs, ABE-packaging RIDE VLPs, and Cas9-packaging VEDIC particles produced from *MAF1*-knockout cells exhibited a 1.7-, 1.8-, and 2.6-fold increase in delivery potency, respectively, compared to the same particles produced from standard Gesicle cells (Extended Data Fig. 5e–g). These results indicate that the improvements conferred by *MAF1*-knockout producer cells are generalizable across several different RNP-packaging cell-derived bioparticles that use distinct particle scaffolds and cargo packaging mechanisms. Taken together, our findings demonstrate that knockout of *MAF1* in producer cells yields consistent improvements in VLP delivery potency across eVLP cargo types, particle types, and producer-cell lines.

### MAF1-knockout producer cells improve sgRNA packaging into eVLPs across different sgRNA expression levels

Finally, we evaluated whether the improvements conferred by *MAF1*-knockout cells could synergize with additional strategies for modulating sgRNA expression levels in producer cells. We observed that increasing the ratio of sgRNA-expressing plasmid to other packaging plasmids transfected during eVLP production did not improve the delivery potency of v4 ABE-eVLPs produced from either standard Gesicle cells or *MAF1*-knockout cells (Extended Data Fig. 6a, b). Increasing the absolute amount of sgRNA-expressing plasmid transfected during eVLP production by up to two-fold also did not improve the delivery potency of v4 ABE-eVLPs produced from either standard Gesicle cells or *MAF1*-knockout cells (Extended Data Fig. 6c). These results indicate that simply increasing the amount of sgRNA-expressing plasmid transfected during eVLP production is not sufficient to improve eVLP potency, possibly because of tradeoffs associated with either reducing the amounts of other packaging plasmids or increasing the total amount of transfected DNA per producer cell.

To further investigate alternative strategies for improving sgRNA expression, we adopted a strategy that was recently developed for EDVs in which additional sgRNA expression cassettes are included in the other packaging plasmids^13^ (Extended Data Fig. 7a). This strategy (hereafter referred to as ++sgRNA) avoids the tradeoffs mentioned above since the amount of sgRNA-encoding DNA is increased without substantially perturbing the amounts of other packaging plasmids or the total amount of transfected DNA per producer cell. We observed that v4 ABE-eVLPs produced from Gesicle cells using ++sgRNA plasmids exhibited a 3.3-fold increase in delivery potency compared to standard v4 ABE-eVLPs produced from Gesicle cells (Fig. 5f). Additionally, we observed that v4 ABE-eVLPs produced from *MAF1*-knockout cells using ++sgRNA plasmids also exhibited a 3.4-fold increase in delivery potency compared to standard v4 ABE-eVLPs produced from *MAF1*-knockout cells (Fig. 5f). These results indicate that the use of ++sgRNA plasmids synergizes with *MAF1*-knockout producer cells to further improve delivery potency. Importantly, v4 ABE-eVLPs produced using ++sgRNA plasmids still exhibited a 2.6-fold increase in delivery potency when produced from *MAF1*-knockout cells compared to Gesicle cells, highlighting the consistent benefit of using *MAF1*-knockout producer cells. Indeed, combining ++sgRNA plasmids and *MAF1*-knockout producer cells yielded the most potent eVLP-mediated base editing at the *BCL11A* enhancer site in HEK293T cells that we observed in this study (Fig. 5f).

We confirmed via RT-qPCR that both the ++sgRNA strategy and knockout of *MAF1* led to increased sgRNA packaging into eVLPs; these increases in sgRNA packaging were additive when both modified strategies were employed (Extended Data Fig. 7b). In particular, we observed that combining both strategies increased the percentage of eVLP-packaged Cas9 molecules complexed with sgRNAs from 20% to 50% (Extended Data Fig. 7b). Collectively, our findings demonstrate that the use of *MAF1*-knockout producer cells can further improve eVLP properties when combined with additional strategies for increasing sgRNA expression during eVLP production.

## Discussion

In this study, we developed a general genome-wide screening approach to comprehensively evaluate how genetic perturbations in producer cells influence eVLP production. Using this approach, we identified the HUSH epigenetic silencing complex and the RNA Pol III repressor MAF1 as key regulators of eVLP protein and sgRNA cargo expression, respectively. Leveraging these insights, we engineered HUSH-knockout producer cells that support a 4-fold increase in eVLP delivery potency when particles are produced using genomically integrated gag–cargo expression, and *MAF1*-knockout producer cells that support a 2- to 9-fold increase in eVLP delivery potency under standard production conditions across several cargo, particle, producer-cell, and target-cell types.

Our results reveal a previously underappreciated role for the HUSH complex in modulating eVLP protein cargo expression in producer cells. Under standard eVLP production conditions, we found that HUSH-knockout producer cells exhibited increased gag–cargo expression, which disrupted the optimal component stoichiometry and reduced eVLP potency. However, under conditions in which gag–cargo expression is limiting, including when particles are produced using genomically integrated gag–cargo expression, we found that the use of HUSH-knockout producer cells improved eVLP delivery potency (Fig. 2f). We therefore anticipate that HUSH-knockout producer cells will be useful for eVLP applications in which genomically integrated gag–cargo expression is required (e.g. barcoded eVLP evolution^22^). Additionally, our results suggest that HUSH-knockout cells are likely to be valuable for future efforts to produce eVLPs or related particles using producer cells that stably express components from genomically integrated cassettes instead of transfected plasmids, a strategy that has proven useful for viral vector manufacturing^27,46–48^.

We anticipate that the use of *MAF1*-knockout cells instead of standard producer cells will advance the utility of VLP delivery in several ways. For one, eVLPs produced from *MAF1*-knockout cells exhibit increased sgRNA packaging and increased delivery potency both in cell culture and in mice compared to standard eVLPs (Fig. 4), suggesting that *MAF1*-knockout producer cells should be used when maximal editing efficiencies are required. Additionally, the gene editing efficiencies at a given dose of eVLPs produced from standard cells can in general be achieved using a 2- to 9-fold lower dose of eVLPs produced from *MAF1*-knockout cells (Fig. 4a–i), suggesting that using *MAF1*-knockout cells for eVLP production will be especially beneficial for applications that are constrained by the maximum administrable eVLP dose. Furthermore, by minimizing the eVLP dose required to achieve efficient editing, the use of *MAF1*-knockout producer cells would also reduce the total volume of producer-cell culture required for eVLP production, which could greatly simplify the application of eVLPs in large-scale studies. Importantly, given that several previous studies have identified sgRNA loading into VLPs and related delivery vehicles as a bottleneck that limits delivery potency^8,13,22,49^, our discovery that *MAF1*-knockout producer cells improve sgRNA loading into VLPs will likely be widely applicable for improving RNP delivery using various cell-derived bioparticles, including the five distinct VLP constructs we tested in this study (Extended Data Fig. 5).

Altogether, our study provides a general workflow for identifying perturbations in producer cells that impact eVLP production and genetically engineering producer cells for improved eVLP production. While previous studies have applied CRISPR screens that use particle-packaged nucleic acid as a way to investigate the cellular determinants of viral vector^24–26^ or extracellular vesicle production^50,51^, our study is the first to our knowledge that leverages insights from a producer-cell screen to improve VLP production and delivery potency. Notably, our results highlight the importance of conducting a producer-cell screen using RNP-packaging VLPs instead of viral vectors, since we uncovered genes that are uniquely relevant for RNP cargo expression and packaging. Additionally, our screen design is highly general and can be applied to evaluate various genetic and epigenetic perturbations on a genome-wide scale, including for CRISPR activation or base editing screens. We anticipate that further improvements in particle production are likely possible by employing varied producer-cell perturbations instead of single gene knockouts alone. Overall, while iterative particle architecture engineering continues to improve VLP delivery potency and cell-type targeting capabilities^10,13,22,52^, our work establishes producer-cell genetic engineering as a distinct and potentially powerful strategy for maximizing the utility of VLP-based delivery modalities.

---

**Extended Data Fig. 1.**
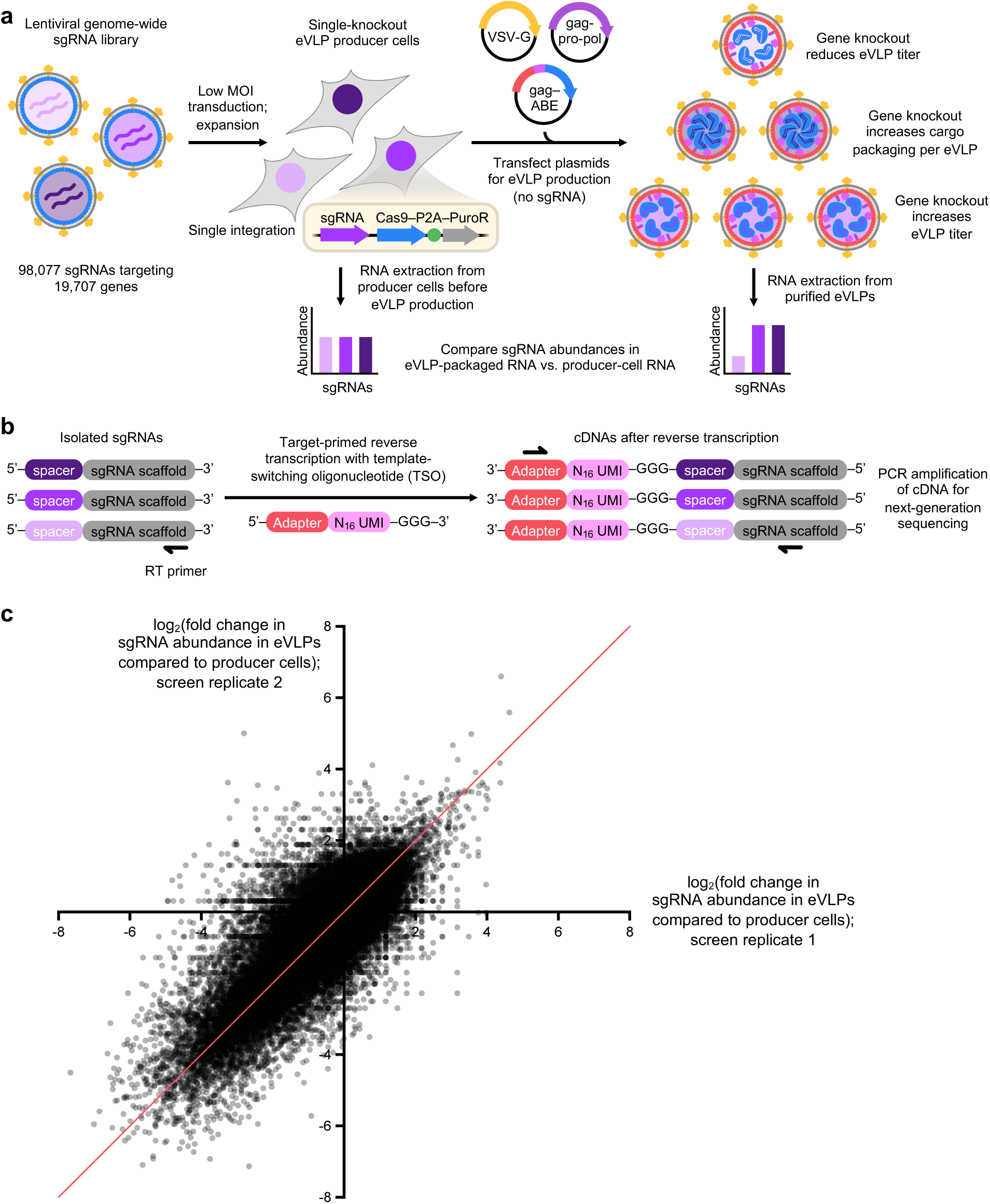
Genome-wide knockout screening for cargo-loaded eVLP production. **a**, Overview of the genome-wide knockout screening approach. Low-MOI Cas9/sgRNA lentiviral transduction was used to introduce a genome-wide sgRNA library into Gesicle producer cells, thereby generating a pool of single-knockout producer cells. eVLP production was then initiated from this pool by transfecting expressing eVLP protein components. No additional sgRNA-encoding plasmid was transfected, ensuring that all eVLPs produced from a given producer cell packaged cargo containing the specific knockout-inducing sgRNA that was already introduced into that producer cell by lentiviral transduction. The abundance of a particular sgRNA in eVLPs compared to producer cells reflects how the corresponding gene knockout influences eVLP production or cargo loading. **b**, Schematic of the template-switching reverse transcription approach used for direct-capture sgRNA spacer sequencing from both eVLP-packaged RNA and producer-cell RNA. UMI=unique molecular identifier. **c**, Fold change in sgRNA abundance in eVLPs compared to producer cells across replicates of the genome-wide knockout screen. Each sgRNA in the library is represented as a single dot and reflects the values from n=2 screen replicates. The y=x line is plotted in red. See also Supplementary Tables 3–5.

**Extended Data Fig. 2.**
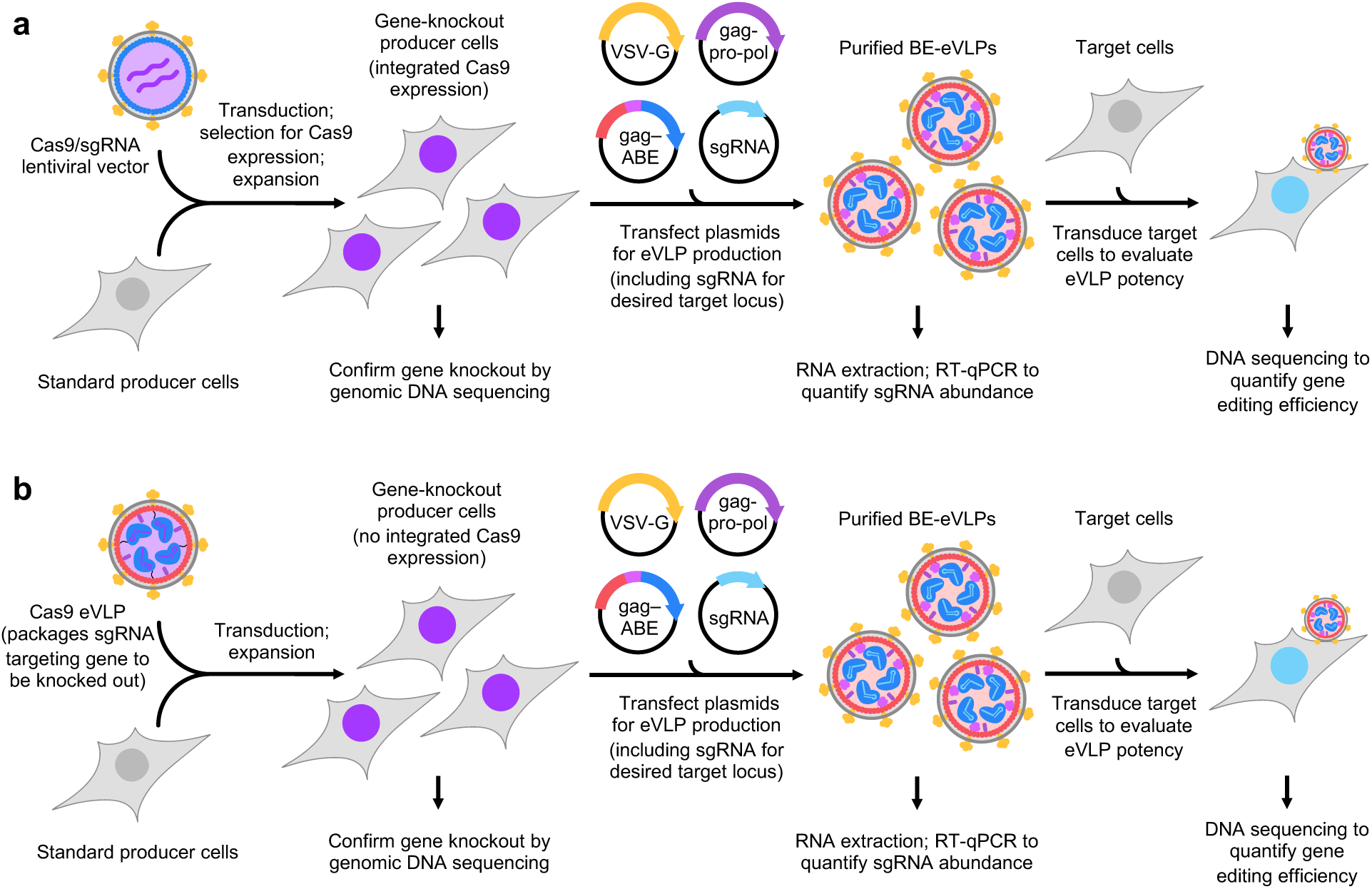
Generating and characterizing single-knockout producer cell lines. **a**, Schematic of generating polyclonal single-knockout producer cells via Cas9/sgRNA lentiviral transduction, followed by characterizing the resulting cell lines and their use for eVLP production. **b**, Schematic of generating polyclonal single-knockout producer cells via v4 Cas9-eVLP transduction, followed by characterizing the resulting cell lines and their use for eVLP production.

**Extended Data Fig. 3.**
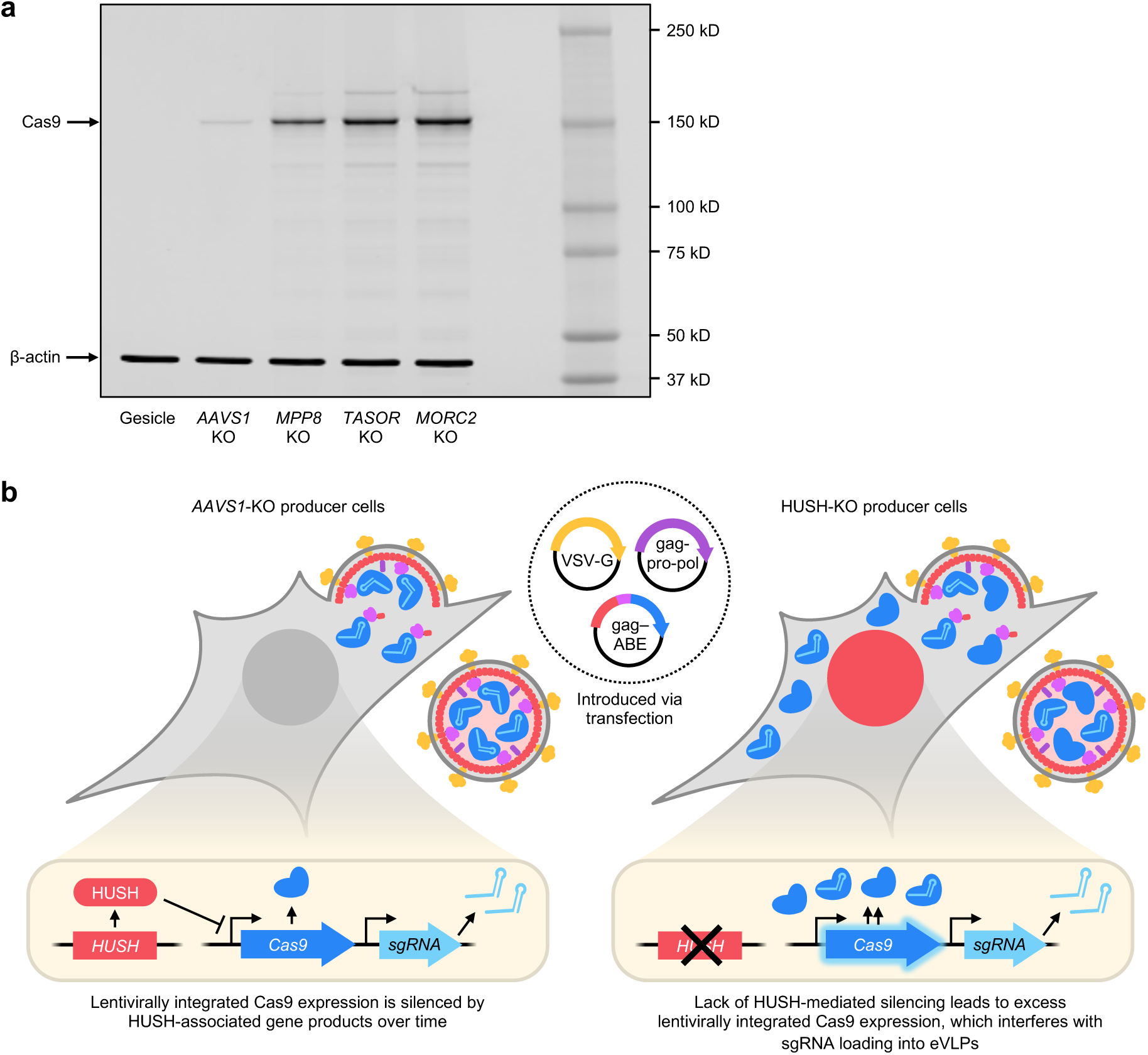
HUSH-knockout producer cells exhibit increased lentiviral Cas9 expression. **a**, Western blot analysis of producer-cell protein lysates. **b**, Proposed model for *AAVS1*-knockout producer cells versus HUSH-knockout producer cells. In *AAVS1*-knockout cells, HUSH-associated gene products silence lentivirally integrated Cas9 expression over time. In HUSH-knockout producer cells, lack of HUSH-mediated silencing leads to excess expression of lentivirally integrated Cas9, which interferes with sgRNA loading into eVLPs and therefore reduces eVLP-packaged sgRNA abundance.

**Extended Data Fig. 4.**
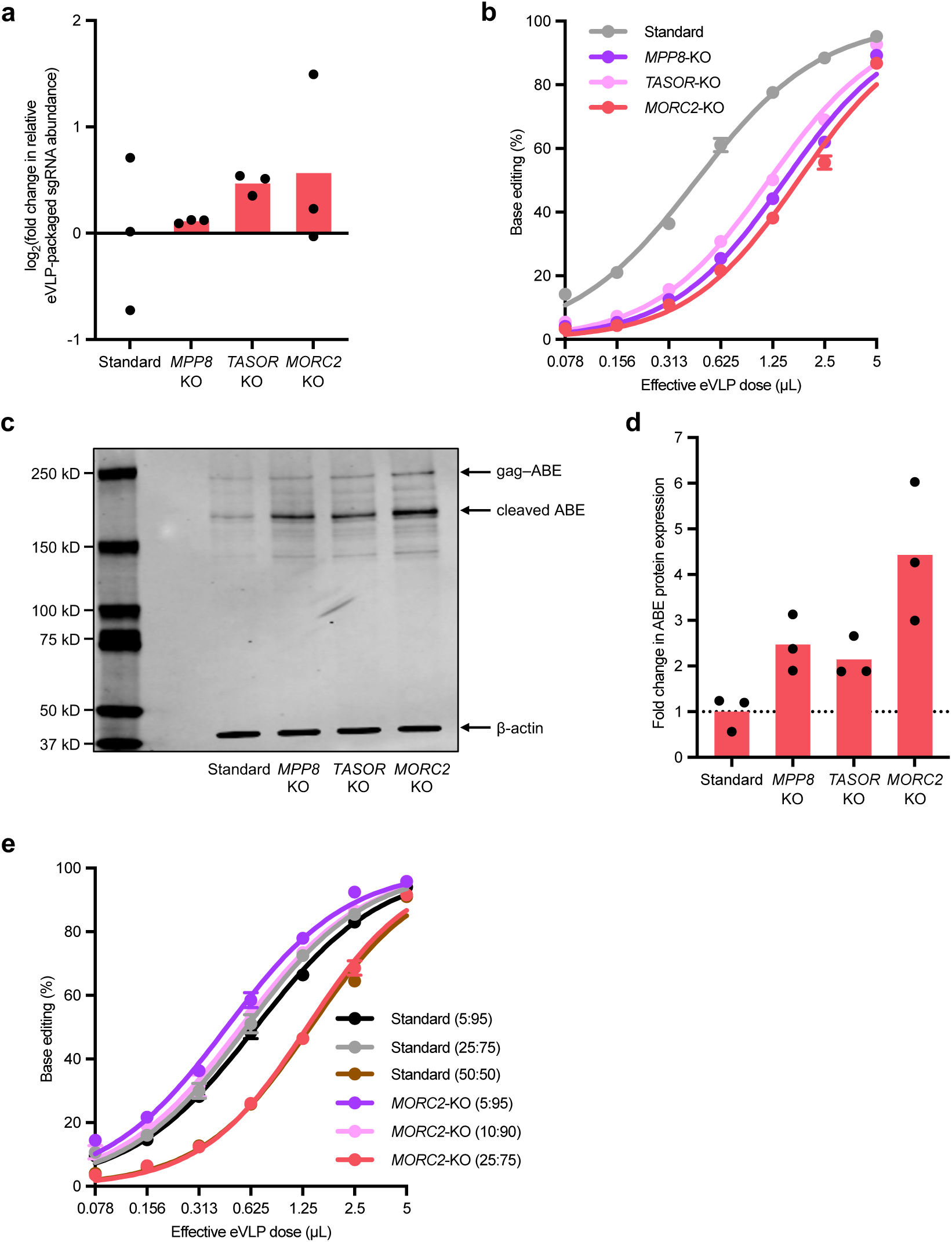
HUSH-knockout producer cells exhibit increased eVLP protein cargo expression. **a**, Fold change in eVLP-packaged sgRNA abundance measured by RT-qPCR, normalized relative to sgRNA abundance in eVLPs produced from Gesicle cells. Bars reflect the mean of n=3 replicates, and dots represent individual replicate values. **b**, Comparison of v4 ABE-eVLPs produced from Gesicle or HUSH-knockout cells across a range of eVLP doses. Adenine base editing efficiencies at position A_7_ of the *BCL11A* +58 enhancer site in HEK293T cells are shown. **c**, Western blot analysis of protein lysates from producer cells transfected with eVLP production plasmids. The full-length gag–ABE fusion is ∼247 kD and the cleaved ABE (following viral protease-mediated cleavage) is ∼184 kD. **d**, Fold change in transfected-producer-cell ABE protein abundance measured by Western blot, normalized relative to ABE protein abundance in transfected Gesicle cells. Bars reflect the mean of n=3 replicates, and dots represent individual replicate values. **e**, Comparison of v4 ABE-eVLPs produced from Gesicle or *MORC2*-knockout cells across a range of eVLP doses using different transfected eVLP plasmid stoichiometries. Values in parentheses reflect the gag–ABE:gag–pro–pol plasmid ratio used for eVLP production. Adenine base editing efficiencies at position A_7_ of the *BCL11A* +58 enhancer site in HEK293T cells are shown. In **b** and **e**, dots and error bars represent mean ± s.d. of n=3 biological replicates. Data were fit to four-parameter logistic curves using nonlinear regression.

**Extended Data Fig. 5.**
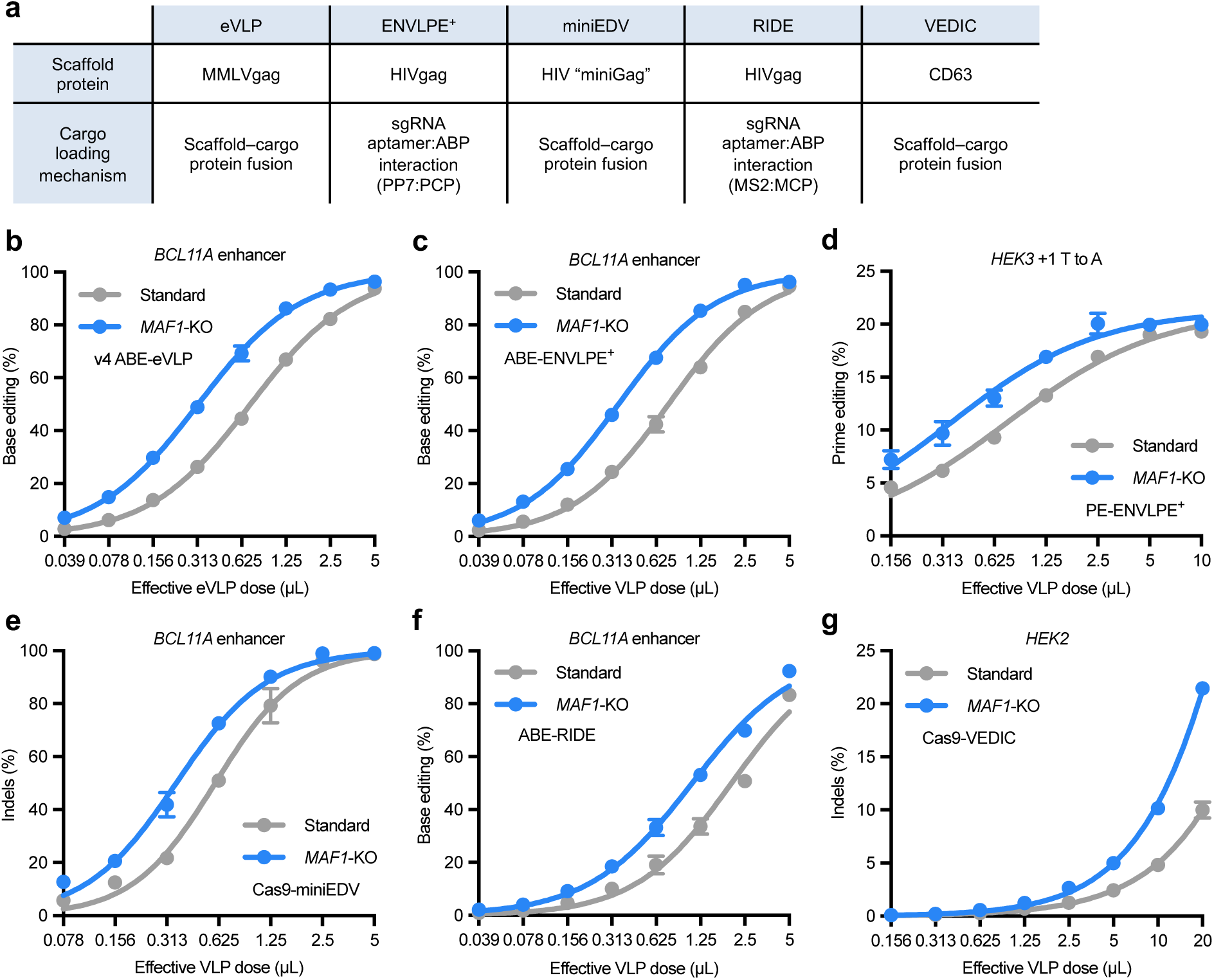
*MAF1*-knockout producer cells improve delivery potency across various particle types. **a**, Summary of RNP-packaging particles that were tested. **b**, Comparison of ABE-packaging eVLPs produced from standard or *MAF1*-knockout Gesicle cells across a range of doses. Adenine base editing efficiencies at position A_7_ of the *BCL11A* +58 enhancer site in HEK293T cells are shown. **c**, Comparison of ABE-packaging ENVLPE^+^ particles produced from standard or *MAF1*-knockout Gesicle cells across a range of doses. Adenine base editing efficiencies at position A_7_ of the *BCL11A* +58 enhancer site in HEK293T cells are shown. **d**, Comparison of PE-packaging ENVLPE^+^ particles produced from standard or *MAF1*-knockout Gesicle cells across a range of doses. +1 T-to-A prime editing efficiencies at the *HEK3* site in HEK293T cells are shown. **e**, Comparison of Cas9-packaging miniEDV particles produced from standard or *MAF1*-knockout Gesicle cells across a range of doses. Indel frequences at the *BCL11A* +58 enhancer site in HEK293T cells are shown. **f**, Comparison of ABE-packaging RIDE particles produced from standard or *MAF1*-knockout Gesicle cells across a range of doses. Adenine base editing efficiencies at position A_7_ of the *BCL11A* +58 enhancer site in HEK293T cells are shown. **g**, Comparison of Cas9-packaging VEDIC particles produced from standard or *MAF1*-knockout Gesicle cells across a range of doses. Indel frequencies at the *HEK2* site in HEK293T cells are shown. **b–g**, Dots and error bars represent mean ± s.d. of n=3 biological replicates. Data were fit to four-parameter logistic curves using nonlinear regression.

**Extended Data Fig. 6.**
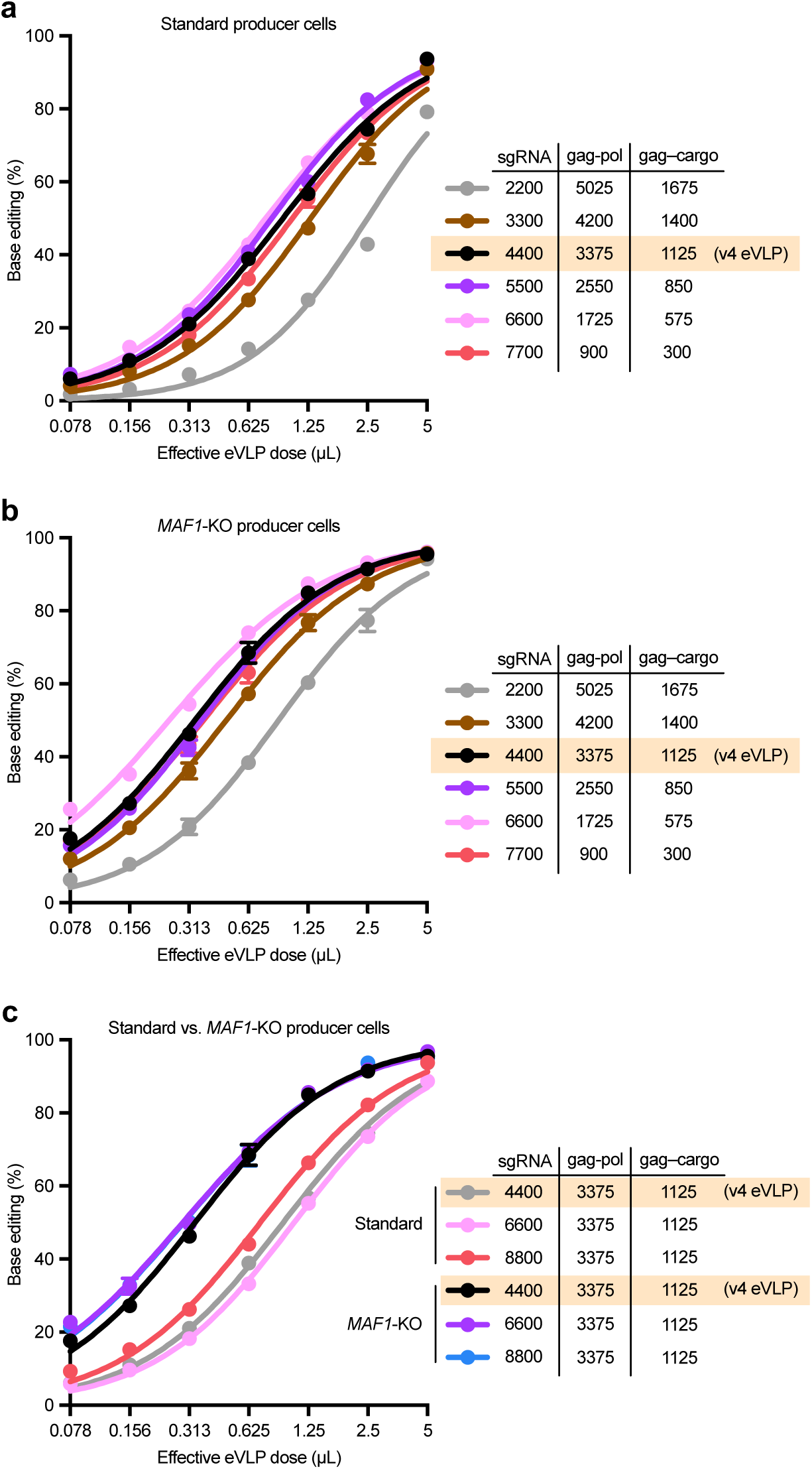
Increasing sgRNA plasmid amounts does not improve eVLP delivery potency. **a–b**, Comparison of v4 ABE-eVLPs produced using different sgRNA plasmid amounts (relative to other packaging components) from standard (**a**) or *MAF1*-knockout (**b**) Gesicle cells across a range of doses. **c**, Comparison of v4 ABE-eVLPs produced using different sgRNA plasmid amounts (supplemented in addition to standard packaging components) from standard or *MAF1*-knockout Gesicle cells across a range of doses. **a–c**, Adenine base editing efficiencies at position A_7_ of the *BCL11A* +58 enhancer site in HEK293T cells are shown. Dots and error bars represent mean ± s.d. of n=3 biological replicates. Data were fit to four-parameter logistic curves using nonlinear regression.

**Extended Data Fig. 7.**
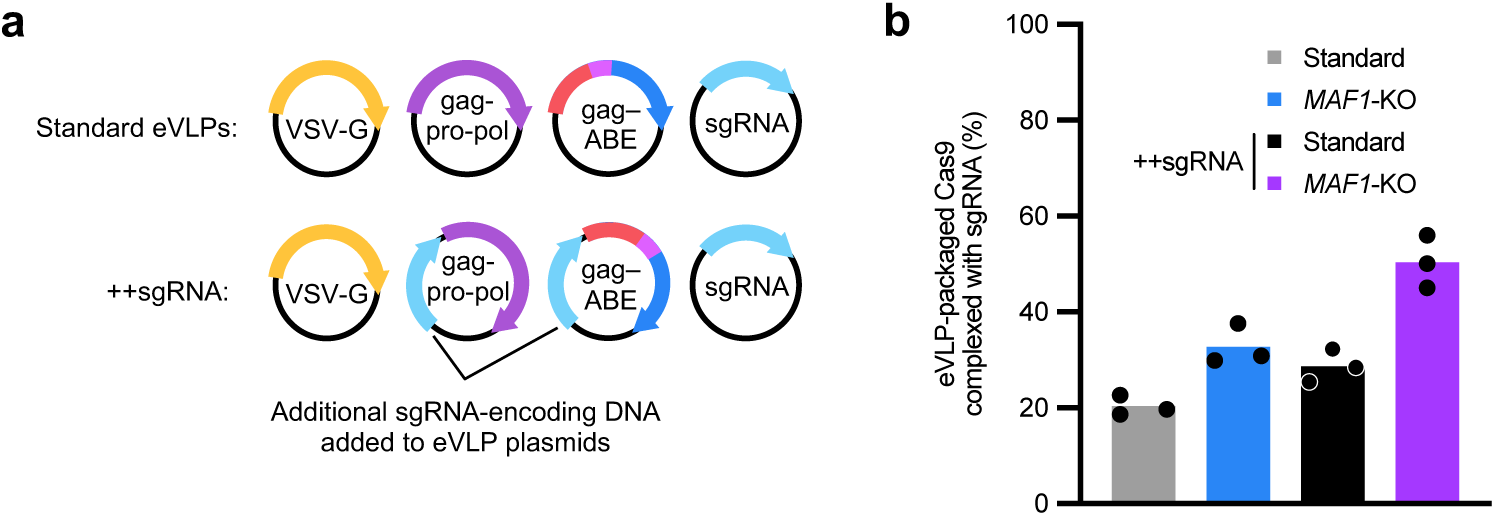
Adding sgRNA expression cassettes to eVLP packaging plasmids improves eVLP sgRNA packaging. **a**, Schematic of additional sgRNA expression cassettes (++sgRNA) added to eVLP packaging plasmids (gag–pro–pol and gag–ABE). **b**, Percentage of eVLP-packaged Cas9 molecules that are complexed with sgRNAs. Absolute sgRNA amounts were quantified by RT-qPCR using *in vitro*-transcribed sgRNA standards, and absolute Cas9 protein amounts were quantified by anti-Cas9 ELISA (see Methods). Bars reflect the mean of n=3 replicates, and dots represent individual replicate values.

## Methods

### Cloning

Plasmids used in this study were cloned via Gibson cloning (NEBuilder^®^ HiFi DNA Assembly; E2621X) or Golden Gate cloning. DNA was amplified via PCR using Phusion HotStart II polymerase (Thermo Fisher Scientific; F537L). Mach1 (Thermo Fisher Scientific; C862003) or NEB^®^ Stable (New England Biolabs; C3040H) chemically competent *E. coli* were used for plasmid propagation. Plasmids were isolated using QIAGEN Plasmid Plus Midi (12945) or Maxi (12963) Kits according to the manufacturer’s protocols.

### Cell culture

HEK293T cells (ATCC; CRL-3216), Gesicle Producer 293T cells (Takara; 632617), HEK293T/17 cells (ATCC; CRL-11268), and Neuro-2a cells (ATCC; CCL-131) were maintained in DMEM + GlutaMAX (Life Technologies; 10569044) supplemented with 10% (v/v) fetal bovine serum (Gibco; A5670701). Cells were cultured at 37 °C with 5% carbon dioxide and were confirmed to be negative for mycoplasma by testing with MycoAlert (Lonza; LT07-318).

### eVLP production and purification

eVLPs were produced as described previously^8,22^. Briefly, eVLPs were produced by transient transfection of Gesicle 293T cells, HEK293T/17 cells, or variants thereof as noted in the main text and figure legends. Producer cells were seeded in T-75 flasks at a density of 5×10^6^ cells per flask in 10 mL of media. After 20–24 h, cells were transfected using the jetPRIME transfection reagent (Polyplus; 114-75) according to the manufacturer’s protocols. For producing eVLPs, a mixture of plasmids expressing VSV-G (400 ng), MMLV gag–pro–pol (3,375 ng), gag–cargo (1,125 ng), and an sgRNA (4,400 ng) was co-transfected per T-75 flask.

40–48 h post-transfection, producer cell supernatant was harvested and centrifuged for 5 min at 500 g to remove cell debris. The clarified eVLP-containing supernatant was filtered through a 0.45-µm PVDF filter (Millipore; SE1M003M00). The filtered supernatant was concentrated 100-fold using PEG-it Virus Precipitation Solution (System Biosciences; LV825A-1) according to the manufacturer’s protocols and resuspended in Opti-MEM serum-free media (Thermo Fisher Scientific; 31985070). For eVLPs prepared for protein or RNA quantification, or those that were injected into mice, the filtered supernatant was concentrated 1500-fold by ultracentrifugation using a cushion of 20% (w/v) sucrose in PBS. Ultracentrifugation was performed at 26,500 rpm (86,242 g) for 2 h (4°C) using an SW32 Ti rotor in an Optima XE Ultracentrifuge (Beckman Coulter). Following ultracentrifugation, eVLP pellets were resuspended in cold PBS pH 7.4 (Gibco; 10010023) and centrifuged at 1,000 g for 5 min to remove debris. eVLPs were frozen at a rate of -1°C/min and stored at -80°C. eVLPs were thawed on ice immediately prior to use.

### eVLP transduction and genomic DNA isolation

Cells were transduced with eVLPs as described previously^8,22^ with a few modifications. Cells were plated for transduction in 96-well plates at a density of 16,000 cells per well. After 20–24 h, eVLPs (that were concentrated 100-fold using PEG precipitation described above) were added directly to the culture media in each well. In almost all cases, eVLPs were serially diluted by 2-fold to generate dose titration curves, and a constant volume of 5 µL of eVLP-containing solution was added to each well. For certain experiments, a constant volume of 10 or 20 µL of VLP-containing solution was added to each well, as noted in the appropriate subfigures. 48–72 h post-transduction, cellular genomic DNA was isolated. Media was aspirated from each well, and 60 µL of lysis buffer (10 mM Tris-HCl pH 8.0, 0.05% SDS, 25 µg mL^−1^ Proteinase K (Thermo Fisher Scientific; EO0492)) was added to each well. Lysis was achieved by incubation at 37 °C for 1–1.5 h followed by heat inactivation at 80°C for 30 min.

### High-throughput sequencing of genomic DNA

Genomic DNA was isolated as described above. Following genomic DNA isolation, 1 µL of the isolated DNA (1–10 ng) was used as input for the first of two PCR reactions. Genomic loci were amplified in PCR1 using Phusion U polymerase (Thermo Fisher Scientific; F562L). PCR1 primers are listed in Supplementary Tables 2 and 7 under the HTS_forward and HTS_reverse columns. PCR1 was performed as follows: 95 °C for 3 min; 30 cycles of 95 °C for 15 s, 61 °C for 20 s, and 72 °C for 30s; 72°C for 1 min. PCR1 products were confirmed on a 1% agarose gel. 1 μL of PCR1 was used as an input for PCR2 to install Illumina barcodes. PCR2 was conducted for nine cycles of amplification using Phusion HotStart II polymerase (Thermo Fisher Scientific; F537L). Following PCR2, samples were pooled and gel purified in a 1% agarose gel using a QIAquick Gel Extraction Kit (Qiagen; 28704). Library concentration was quantified using the Qubit^TM^ dsDNA High-Sensitivity Assay Kit (Thermo Fisher Scientific; Q33230). Samples were sequenced on an Illumina MiSeq instrument (paired-end read, read 1: 200–280 cycles, read 2: 0 cycles) using an Illumina MiSeq 300 v2 Kit (Illumina).

### High-throughput sequencing data analysis for gene editing experiments

Sequencing reads were demultiplexed using the MiSeq Reporter software (v2.6) (Illumina) and were analyzed using CRISPResso2 (ref. ^53^) (v2.3.2) as previously described^8^. Batch analysis mode (one batch for each unique amplicon and sgRNA combination analyzed) was used in all cases. Reads were filtered by minimum average quality score (Q > 30) prior to analysis. For analysis of base editing efficiencies, the following quantification window parameters were used: -w 20 -wc -10. Base editing efficiencies are reported as the percentage of sequencing reads containing a given base conversion at a specific position. For analysis of prime editing efficiencies, the following quantification parameters were used: -w 30 -wc -3 –dicard_indel_reads. Prime editing efficiencies are reported as the number of reads aligned to the edited amplicon sequence divided by the number of reads aligned to all amplicons multiplied by 100. Prism 10 (GraphPad) was used to generate dot plots and bar plots. Amplicon sequences are provided in Supplementary Tables 2 and 7.

### Lentiviral vector production

Lentiviral vectors were constructed via Golden Gate cloning into the LentiCRISPRv2-Opti backbone, a gift from the Whitehead Institute Functional Genomics Platform (Addgene #163126). Plasmids were propagated in NEB^®^ Stable (New England Biolabs; C3040H) chemically competent *E. coli*. Prior to lentiviral vector production, HEK293T/17 cells were maintained in antibiotic-free DMEM supplemented with 10% fetal bovine serum (v/v). On day 1, 5×10^6^ cells were plated in 10 mL of media in T-75 flasks. The following day, cells were transfected with 6 µg of VSV-G envelope plasmid, 9 µg of psPAX2 (plasmid encoding viral packaging proteins) and 9 µg of transfer vector plasmid diluted in 1,500 µL Opti-MEM with 70 µL of FuGENE HD transfection reagent (Promega; E2312). Two days after transfection, media was centrifuged at 500 *g* for 5 min to remove cell debris followed by filtration using a 0.45-µm PVDF vacuum filter (Millipore; SE1M003M00). The filtered supernatant was concentrated using PEG-it Virus Precipitation Solution (System Biosciences; LV825A-1) according to the manufacturer’s protocols and resuspended in Opti-MEM serum-free media (Gibco; 31985070).

### sgRNA library propagation

For the genome-wide knockout screen, a previously described human genome-wide sgRNA library containing 98,077 unique sgRNAs targeting 19,707 genes across the human genome^30^ was obtained as a gift from the Whitehead Institute Functional Genomics Platform. sgRNA sequences are listed in Supplementary Table 3. Vectors contained pU6-sgRNA expression cassettes along with pEFS-Cas9-P2A-PuroR (LentiCRISPRv2-Opti backbone, Addgene #163126). The plasmid library was propagated as follows. First, electrocompetent cells were generated from NEB^®^ Stable (New England Biolabs; C3040H) chemically competent *E. coli* by growing single colonies to mid-log phase in LB media containing 10 µg/mL tetracycline and 50 µg/mL streptomycin, collecting cells by centrifugation at 5,000 g for 1 min at 4 °C, washing with cold 10% (v/v) glycerol, and repeating for a total of four washes. Freshly prepared electrocompetent cells were transferred to a chilled 0.1 cm electroporation cuvette (Bio-Rad; 1652089) and mixed with the purified, assembled library plasmids. Cells were electroporated using a time-constant protocol with t = 5 ms at 1.5 kV. Electroporated cells were recovered at 37 °C for 25 min with shaking. Recovered cells were plated onto 500 cm^2^ plates containing LB media + 1.5% agar supplemented with 100 µg/mL carbenicillin and incubated for 16 h at 37 °C. After overnight incubation, colonies were scraped into LB media and cells were collected by centrifugation. Plasmids were purified using a Plasmid Plus Maxi Kit (Qiagen; 12963) according to the manufacturer’s protocols. sgRNA representation in the propagated plasmid library was confirmed via targeted amplicon sequencing.

### Genome-wide knockout screen for cargo-loaded eVLP production

Lentiviral libraries were produced as described above and were concentrated ∼66-fold by PEG precipitation. Gesicle cells were seeded at a density of 1.5×10^7^ cells per T-175 and immediately transduced with 105 µL of concentrated lentivirus. A total of 2.4×10^8^ Gesicle cells were seeded for transduction. 24 h after transduction, cells were trypsinized and reseeded at half the original density in media supplemented with 2 µg/mL puromycin. Puromycin selection was continued for 7 days, during which time cell viability was monitored and cells were expanded upon reaching confluency. The initial multiplicity of infection (MOI) was inferred by counting surviving cells and assuming a doubling time of 24 h. Using this method, the MOI was calculated to be 0.28, corresponding to approximately 660x coverage of the sgRNA library. A minimum of 1×10^8^ cells were carried forward each time the cells were passaged to retain overall coverage.

After cells were robustly surviving puromycin selection (∼1 week), cells were seeded for eVLP production in T-75 flasks at a density of 5×10^6^ cells/flask. At this time, 1×10^8^ cells were collected by centrifugation, and the cell pellet was frozen at -80°C and reserved for RNA extraction (see below). 24 h after seeding, cells were transfected using the jetPRIME transfection reagent (Polyplus; 114-75) according to the manufacturer’s protocols. A mixture of pUC19 (4,400 ng), VSV-G (400 ng), MMLV gag–pro–pol (3,375 ng), and MMLVgag–ABE8e (1,125) plasmids was co-transfected per T-75 flask. 42 h post transfection, eVLPs were harvested and filtered as described above. The filtered supernatant was concentrated by sucrose-cushion ultracentrifugation as described above. Following ultracentrifugation, eVLP pellets were resuspended in cold PBS pH 7.4 (Gibco; 10010023). Purified eVLPs were treated with DNase (Qiagen; 79254) followed by RNA extraction using the QIAmp Viral RNA Mini Kit (Qiagen; 52904) according to the manufacturer’s protocols. RNA was extracted from the producer-cell pellet using QIAzol Lysis Reagent (Qiagen; 79306) according to the manufacturer’s protocols. This entire process of seeding single-knockout Gesicle cells followed by eVLP production and RNA extractions was performed twice on different days for two screen replicates.

Following RNA extraction, both producer-cell and eVLP RNA were reverse transcribed using the Maxima Reverse Transcriptase (Thermo Fisher Scientific; EP0742) according to the manufacturer’s protocols. Primers and other oligonucleotides used for target-primed sgRNA reverse transcription are listed in Supplementary Table 1. The RT primer was added at a final concentration of 1 µM in the RT reaction. Following incubation at 50°C for 20 min, the template-switching oligo (TSO) was spiked in to the reaction at a final concentration of 1 µM. The reaction was incubated for a further 20 min at 50°C, and heat inactivated at 85°C for 5 min. For producer-cell RNA only, the RT products were further processed by hydrolyzing the RNA in 0.2 M NaOH (final concentration) and heating at 90°C for 10 min, cleaning up with the MinElute Reaction Cleanup Kit (Qiagen; 28104) according to the manufacturer’s protocols, and left size selection using SPRIselect (Beckman Coulter; B23317) at a 1.5x bead ratio to remove smaller products and residual primers. Producer-cell cDNAs were subsequently amplified for sequencing with the KAPA HiFi HotStart ReadyMix (Roche Diagnostics; 09420398001) using 2.5 µL of cDNA input per 25 µL reaction and the following conditions: 95 °C for 3 min; 20 cycles of 98 °C for 20 s, 61 °C for 20 s, and 72 °C for 30s; 72°C for 2 min. eVLP cDNAs were amplified for sequencing with the KAPA HiFi HotStart ReadyMix using 2.5 µL of cDNA input per 25 µL reaction and the following conditions: 95 °C for 3 min; 14 cycles of 98 °C for 20 s, 61 °C for 20 s, and 72 °C for 30s; 72°C for 2 min. Illumina barcodes were installed using KAPA HiFi HotStart ReadyMix using 2 µL of PCR1 input per 25 µL reaction and the following conditions: 95 °C for 3 min; 9 cycles of 98 °C for 20 s, 61 °C for 20 s, and 72 °C for 30s; 72°C for 2 min. Following PCR2, samples were gel purified in a 1% agarose gel using a QIAquick Gel Extraction Kit (Qiagen; 28704). Library concentration was quantified using the Qubit High-Sensitivity Assay Kit (Thermo Fisher Scientific; Q33230). Samples were sequenced on an Illumina NextSeq instrument (paired-end read, read 1: 76 cycles, read 2: 0 cycles) using an Illumina NextSeq 75-cycle High-Output v2.5 Kit (Illumina).

### Genome-wide knockout screen analysis

Code used for processing sequencing reads from the genome-wide knockout screen is provided in Supplementary Note 1. Briefly, sequencing reads were demultiplexed using bcl2fastq (v2.20.0.422). Reads were trimmed using seqkit^54^ (v2.6.1) to restrict analysis to the UMI and sgRNA protospacer sequences only. Then, reads were UMI-deduplicated using AmpUMI^55^ and trimmed with seqkit to remove the UMI and restrict subsequent analysis to the sgRNA protospacer sequences only. Processed reads were aligned to a reference fasta file containing all sgRNA sequences using bowtie^56^ (v1.3.1), and sgRNA counts were generated from alignments as previously described^30^. Raw sgRNA counts are provided in Supplementary Table 4. MAGeCK^57^ (v0.5.9.5) was used to quantify gene-level phenotypes from the sgRNA counts. Producer-cell samples were treated as “control” samples and eVLP samples were treated as “treatment” samples for the purpose of MAGeCK analysis. Median-normalized sgRNA read counts and fold changes are provided in Supplementary Table 5, and gene-level phenotypes are provided in Supplementary Table 6. For genes with a positive median log_2_ fold change, the positive MAGeCK RRA score, p-value, and FDR were used, and for genes with a negative median log_2_ fold change, the negative MAGeCK RRA score, p-value, and FDR were used. Prism 10 (GraphPad) was used to generate dot plots.

### Generation of knockout cell lines

To generate knockout cell lines via Cas9/sgRNA lentiviral transduction, lentiviral vectors were cloned and produced as described above. Cells were seeded in T-75 flasks at a density of 5×10^6^ cells/flask and immediately transduced with 50 µL of concentrated lentivirus. 24 h after transduction, cells were trypsinized and reseeded in media supplemented with 2 µg/mL puromycin. Puromycin selection was continued, during which time cell viability was monitored and cells were expanded upon reaching confluency. After cells were robustly surviving puromycin selection (∼1 week), cells were harvested and genotyped via targeted-amplicon high-throughput sequencing as described above to confirm gene knockout. Cell lines were regularly genotyped over time to confirm continued gene knockout.

To generate knockout cell lines via Cas9-eVLP transduction, v4 Cas9-eVLPs were produced as described above. Cells were seeded in 96-well plates at a density of 16,000 cells per well. 24 h after seeding, cells were transduced with 5 µL of purified Cas9-eVLPs per well. 72 h after eVLP transduction, cells were harvested and genotyped via targeted-amplicon high-throughput sequencing as described above to confirm gene knockout. Cells were expanded and regularly genotyped over time to confirm continued gene knockout.

### eVLP-packaged sgRNA extraction and RT-qPCR

eVLPs for RNA extraction were purified via sucrose-cushion ultracentrifugation as described above. Purified eVLPs were treated with DNase (Qiagen; 79254) followed by RNA extraction using the QIAmp Viral RNA Mini Kit (Qiagen; 52904) according to the manufacturer’s protocols. Extracted RNA was reverse transcribed using SuperScript^TM^ III First-Strand Synthesis SuperMix (Thermo Fisher Scientific; 18080400) and an sgRNA-specific DNA primer (see Supplementary Table 1) according to the manufacturer’s protocols. qPCR analysis of the resulting cDNA was performed using a QuantStudio^TM^ 3 Real-Time PCR System (Thermo Fisher Scientific) with SYBR green dye (Lonza; 50512). The relative eVLP-packaged sgRNA abundance was calculated as log_2_[fold change] (ΔC_t_) relative to control eVLPs (i.e. those produced from standard or *AAVS1*-knockout cells).

### Producer-cell RNA extraction and RT-qPCR

Cellular RNA was extracted using the RNeasy Plus Mini Kit (Qiagen; 74136) according to the manufacturer’s protocols. Extracted RNA was reverse transcribed using SuperScript^TM^ III First-Strand Synthesis SuperMix (Thermo Fisher Scientific; 18080400) using random hexamer primers according to the manufacturer’s protocols. qPCR analysis of the resulting cDNA was performed using a QuantStudio^TM^ 3 Real-Time PCR System (Thermo Fisher Scientific) with SYBR green dye (Lonza; 50512). The relative Cas9 mRNA abundance across different producer cells was calculated via the ΔΔC_t_ method using *ACTB* as the housekeeping gene. qPCR primers are listed in Supplementary Table 1.

### Producer-cell protein extraction and Western blot analyses

For producer-cell protein extraction, 1×10^6^ cells were collected by centrifugation at 500 g for 5 min. Cell pellets were washed once with 1 mL of cold PBS and lysed using RIPA buffer (Thermo Fisher Scientific; 89901) supplemented with cOmplete^TM^ protease inhibitor cocktail (Roche Diagnostics; 11836153001) according to the manufacturer’s protocols. Cells were lysed at 4°C for 30 min with gentle agitation. The lysate was clarified by centrifugation at 10,000 g for 20 min at 4°C, and the clarified supernatant was combined with 4x Laemmli Sample Buffer (Bio-Rad; 1610747) and heated at 95°C for 10 min. Protein extracts were subsequently separated by electrophoresis at 150 V for 45 min on a NuPAGE 3–8% Tris-Acetate gel (Thermo Fisher Scientific; EA0378BOX) in 1x NuPAGE Tris-Acetate SDS running buffer (Thermo Fisher Scientific; LA0041). Transfer to a PVDF membrane was performed using an iBlot 2 Gel Transfer Device (Thermo Fisher Scientific) at 20 V for 7 min. The membrane was blocked for 1 h at room temperature with rocking in blocking buffer: 1% bovine serum albumin (BSA) and 0.1% Tween-20 in TBS (150 mM NaCl and 50 mM Tris-HCl). After blocking, the membrane was incubated overnight at 4°C with rocking with mouse anti-Cas9 (Cell Signaling Technology; 14697, 1:1000 dilution) and rabbit anti-actin (Cell Signaling Technology; 4970, 1:1000 dilution). The membrane was washed three times with 1xTBST (150 mM NaCl, 0.5% Tween-20, and 50 mM Tris-HCl) for 10 min each time at room temperature, then incubated with goat anti-mouse secondary antibody (LI-COR IRDye 680RD; 926-68070, 1:10000 dilution in blocking buffer) and goat anti-rabbit secondary antibody (LI-COR IRDye 800CW; 926-32211, 1:10000 dilution in blocking buffer) for 1 h at room temperature with rocking. The membrane was washed as before and imaged using an Odyssey Imaging System (LI-COR). Protein abundances were quantified by densitometry using ImageJ (v2.14.0). Cas9 abundances were normalized to the actin abundance within each sample, and comparisons across samples were performed using these actin-normalized values.

For protein extraction from cells transfected with eVLP production plasmids, cells were first seeded for eVLP production in T-75 flasks as described above. 24 h after seeding, cells were transfected with all eVLP production plasmids except the VSV-G plasmid. 48 h after transfection, cells were trypsinized and collected by centrifugation, and were then processed as described above.

### eVLP protein content quantification

eVLP protein content quantification was performed as described previously^8^. Briefly, ultracentrifuge-purified eVLPs were lysed in dye-free Laemmli sample buffer (50 mM Tris-HCl pH 7.0, 2% sodium dodecyl sulfate (SDS), 10% (v/v) glycerol, 2 mM dithiothreitol (DTT)) by heating at 95°C for 15 min. The concentration of ABE protein in purified eVLPs was quantified using the FastScan^TM^ Cas9 (*S. pyogenes*) ELISA kit (Cell Signaling Technology; 29666C) according to the manufacturer’s protocols. Recombinant Cas9 (*S. pyogenes*) nuclease protein (New England Biolabs; M0386) was used to generate the standard curve for quantification. The concentration of MLV p30 protein in purified eVLPs was quantified using the MuLV Core Antigen ELISA kit (Cell Biolabs; VPK-156) according to the manufacturer’s protocols. To calculate eVLP titer in particles/µL, the p30 concentration in ng/mL determined by ELISA was multiplied by a factor 2030.57, which accounts for molarity conversions, assumes that 20% of p30 molecules in solution are associated with particles, and assumes a copy number of 1800 molecules of p30 per eVLP, as previously described^8^. The number of ABE protein molecules per eVLP was calculated by determining the ratio of Cas9 molecules to p30 molecules and assuming a copy number of 1800 molecules of p30 per eVLP, as previously described^8^.

### Animal experiments

All mouse experiments were compliant with relevant ethical regulations and were approved by the MIT Committee on Animal Care (D16-00078; 2402000629). Wild-type 8-week-old adult male C57BL/6J mice (000664) were purchased from the Jackson Laboratory. All mice were housed in a room maintained on a 12 h light and dark cycle with *ad libitum* access to standard rodent diet and water. Animals were randomly assigned to various experimental groups.

Retro-orbital injections were performed as described previously^8^ with a few modifications. eVLPs for retro-orbital injections were purified via sucrose-cushion ultracentrifugation as described above. Immediately prior to injection, eVLPs were partially clarified by centrifugation at 1000 g for 5 min to remove debris. The eVLP-containing supernatant was diluted with 0.9% NaCl (Hospira; 00409-4888-10), and a total of 100 µL solution was injected per mouse: 85 µL saline + 15 µL of 1500-fold concentrated purified eVLPs, containing 3×10^11^ eVLPs as was quantified by ELISA using the procedure described above). Genomic DNA was purified from bulk liver tissue using the Agencourt DNAdvance kit (Beckman Coulter; A48705) according to the manufacturer’s protocols. Purified genomic DNA was amplified for high-throughput sequencing as described above, with 25 cycles of amplification during PCR1.

### Production of other VLP constructs

ENVLPE^+^ particles^10^, miniEDVs^44^, RIDE VLPs^16^, and VEDIC^45^ particles were produced analogously to eVLPs. Producer cells were seeded in T-75 flasks at a density of 5×10^6^ cells per flask. After 20–24 h, cells were transfected using the jetPRIME transfection reagent (Polyplus; 114-75) according to the manufacturer’s protocols.

For producing ABE-ENVLPE^+^ particles, a mixture of plasmids expressing VSV-G (1,550 ng), psPAX2-D64V (1,149 ng; Addgene #63586, a gift from David Rawlings & Andrew Scharenberg), pCMV_ENVLPE+ (665 ng; Addgene #232427, a gift from Gil Westmeyer), pCMV_ABE8e_superNLS (1,023 ng; Addgene #232429, a gift from Gil Westmeyer), and a *BCL11A*-targeting PP7-containing sgRNA (4,913 ng; cloned from Addgene #232432, a gift from Gil Westmeyer) was co-transfected per T-75 flask.

For producing PE-ENVLPE^+^ particles, a mixture of plasmids expressing VSV-G (1,550 ng), psPAX2-D64V (1,149 ng; Addgene #63586, a gift from David Rawlings & Andrew Scharenberg), pCMV_ENVLPE+ (665 ng; Addgene #232427, a gift from Gil Westmeyer), pCMV_iPE-C_P2A_Csy4 (1,023 ng; Addgene #232428, a gift from Gil Westmeyer), and pU6_HEK3+1T>A_(PP7-C4-Q1) (4,913 ng; Addgene #232437, a gift from Gil Westmeyer) was co-transfected per T-75 flask.

For producing miniEDVs, a mixture of plasmids expressing VSV-G (930 ng), a *BCL11A*-targeting sgRNA and miniGag (2,790 ng; cloned from Addgene #228957, a gift from Jennifer Doudna), and a *BCL11A*-targeting sgRNA and miniGag–Cas9 (5,580 ng; cloned from Addgene #228958, a gift from Jennifer Doudna) was co-transfected per T-75 flask.

For producing RIDE VLPs, a mixture of plasmids expressing VSV-G (760 ng), pRSV-Rev (610 ng; Addgene #12253, a gift from Didier Trono), pMS2M-PH-gagpol-D64V (1,320 ng; Addgene #166031, a gift from Yujia Cai), pMDLg/pRRE (1,320 ng; Addgene #12251, a gift from Didier Trono), ABE8e (2,640 ng; Addgene #138489, a gift from David Liu), and a *BCL11A*-targeting MS2-containing sgRNA (2,640; Addgene #229772, a gift from Yujia Cai) was co-transfected per T-75 flask.

For producing VEDIC particles, a mixture of plasmids expressing VSV-G (400 ng), a CD63–intein–Cas9 fusion (3,500 ng), and *HEK2*-targeting sgRNA (5,400 ng) was co-transfected per T-75 flask.

For all VLPs, 40–48 h post-transfection, producer cell supernatant was harvested and centrifuged for 5 min at 500 g to remove cell debris. The clarified VLP-containing supernatant was filtered through a 0.45-µm PVDF filter (Millipore; SE1M003M00). The filtered supernatant was concentrated 100-fold using PEG-it Virus Precipitation Solution (System Biosciences; LV825A-1) according to the manufacturer’s protocols and resuspended in Opti-MEM serum-free media (Thermo Fisher Scientific; 31985070). For all VLPs, transduction of HEK293T cells was performed as described above for eVLPs.

### Quantification of eVLP-packaged Cas9 molecules complexed with sgRNAs

For each construct, the number of Cas9 molecules per unit volume of eVLPs was calculated via anti-Cas9 ELISA as described above. For sgRNA quantification, purified eVLPs were treated with DNase (Qiagen; 79254). Then, 100 ng of 151-bp spike-in RNA (see below) was added to each sample, followed by RNA extraction using the QIAmp Viral RNA Mini Kit (Qiagen; 52904) according to the manufacturer’s protocols. Extracted RNA was reverse transcribed using SuperScript^TM^ III First-Strand Synthesis SuperMix (Thermo Fisher Scientific; 18080400) and both an sgRNA-specific DNA primer and a spike-in RNA-specific DNA primer (see Supplementary Table 1) according to the manufacturer’s protocols. qPCR analysis of the resulting cDNA was performed with both sgRNA-specific and spike-in RNA-specific primers (see Supplementary Table 1) using a QuantStudio^TM^ 3 Real-Time PCR System (Thermo Fisher Scientific) with SYBR green dye (Lonza; 50512).

To determine the absolute amount of sgRNA and spike-in cDNA in each sample, a standard curve for each cDNA species was generated. First, RNA standards for both species were generated via *in vitro* transcription using the HiScribe^®^ T7 Quick High Yield RNA Synthesis Kit (New England Biolabs; E2050S). Transcribed RNA yield was quantified using the Qubit^TM^ RNA High-Sensitivity Assay Kit (Thermo Fisher Scientific; Q32852). Then, serial dilutions of each RNA standard were made in water, and the resulting diluted RNAs were subjected to reverse transcription using either sgRNA-specific or spike-in RNA-specific DNA primers as described above. The resulting cDNAs were used as input into qPCR reactions to generate a standard curve for each cDNA species.

Standard curves were analyzed via linear regression using Prism 10 (GraphPad), and these curves were used to calculate the number of sgRNA and spike-in RNA molecules present in each eVLP-derived sample after RNA extraction. To determine the number of sgRNA molecules present in each eVLP sample before RNA extraction, RNA loss during the extraction process was accounted for by calculating the percent recovery of spike-in RNA for each sample (using the knowledge that exactly 100 ng of spike-in RNA was added to each sample before RNA extraction, as described above). Using this RNA recovery factor, the number of sgRNA molecules present in each eVLP sample before RNA extraction was calculated (with the assumption that, for each sample, the percent recovery of spike-in RNA and sgRNA are equal since both species are of similar size). Finally, the percentage of eVLP-packaged Cas9 molecules complexed with sgRNAs was calculated for each sample by dividing the number of sgRNA molecules per unit volume of eVLPs by the number of Cas9 molecules per unit volume of eVLPs. This calculation assumes that all sgRNA molecules detected in eVLPs are complexed with Cas9 molecules, since sgRNA packaging into eVLPs in the absence of Cas9 is negligible^22^.

### Statistical analysis

Data are presented as mean and standard deviation (s.d.). No statistical methods were used to predetermine sample size. Statistical analysis was performed using Prism 10 (GraphPad). Sample sizes are described in the figure legends.

## Data availability

The high-throughput sequencing data generated during this study is being deposited at the NCBI Sequence Read Archive database under PRJNA1288558. Raw data from the genome-wide knockout screen is provided in Supplementary Tables 3–6. Key plasmids from this work will be deposited to Addgene for distribution. Other plasmids and cell lines are available from the corresponding author on request.

## Code availability

The code used for analyzing base editing efficiencies are available at https://github.com/pinellolab/CRISPResso2. The code used for analyzing CRISPR screen results are available in Supplementary Note 1 and at https://sourceforge.net/p/mageck/wiki/Home/.

## Acknowledgements

This work was supported by the Whitehead Institute and Whitehead Fellows Program (A.R.); NIH Director’s Early Independence Award DP5OD037342 (A.R.); Merkin Institute for Transformative Technologies in Healthcare (A.R.); McGuire Family Foundation (A.R.); James M. and Cathleen D. Stone Foundation (A.R.); Owens Family Foundation (A.R.); and Valhalla Foundation (A.R.). H.J. acknowledges support from the National Research Foundation of Korea. A.S. was supported by the Whitehead Scholars Program at Williams College. We thank H. Keys and the Functional Genomics Platform at Whitehead Institute for providing the genome-wide sgRNA library used in our study; A. Nelson, D. Liu, and the Whitehead Genome Technology Core for access to Illumina sequencing instrumentation; and the Jain, Reddien, Weissman, and Young Labs for generous access to equipment.

## Author contributions

A.R. conceived the project. D.L., H.J., A.G., A.S., and A.R. designed the research, performed experiments, and analyzed data. H.J. performed mouse experiments. A.R. wrote the manuscript, and all authors edited the manuscript.

## Competing interests

The authors declare competing financial interests: D.L. and A.R. have filed a patent application on this work through the Whitehead Institute. The remaining authors declare no competing interests.

## Notes

### Summary of Updates

New data added; manuscript text revised; figures revised.

## References

1. Raguram, A., Banskota, S. & Liu, D.R. Therapeutic in vivo delivery of gene editing agents. Cell 185, 2806–2827 (2022).

2. van Haasteren, J., Li, J., Scheideler, O.J., Murthy, N. & Schaffer, D.V. The delivery challenge: fulfilling the promise of therapeutic genome editing. Nat Biotechnol 38, 845–855 (2020).

3. Wang, D., Zhang, F. & Gao, G. CRISPR-Based Therapeutic Genome Editing: Strategies and In Vivo Delivery by AAV Vectors. Cell 181, 136–150 (2020).

4. Wei, T. et al. Delivery of Tissue-Targeted Scalpels: Opportunities and Challenges for In Vivo CRISPR/Cas-Based Genome Editing. ACS Nano 14, 9243–9262 (2020).

5. Wang, J.Y. & Doudna, J.A. CRISPR technology: A decade of genome editing is only the beginning. Science 379, eadd8643 (2023).

6. Tsuchida, C.A., Wasko, K.M., Hamilton, J.R. & Doudna, J.A. Targeted nonviral delivery of genome editors in vivo. Proc Natl Acad Sci U S A 121, e2307796121 (2024).

7. An, M., et al. Engineered virus-like particles for transient delivery of prime editor ribonucleoprotein complexes in vivo. Nat Biotechnol 42, 1526–1537 (2024).

8. Banskota, S. et al. Engineered virus-like particles for efficient in vivo delivery of therapeutic proteins. Cell 185, 250–265 e216 (2022).

9. Choi, J.G. et al. Lentivirus pre-packed with Cas9 protein for safer gene editing. Gene Ther 23, 627–633 (2016).

10. Geilenkeuser, J., et al. Engineered nucleocytosolic vehicles for loading of programmable editors. Cell 188, 2637–2655 e2631 (2025).

11. Haldrup, J. et al. Engineered lentivirus-derived nanoparticles (LVNPs) for delivery of CRISPR/Cas ribonucleoprotein complexes supporting base editing, prime editing and in vivo gene modification. Nucleic Acids Res 51, 10059–10074 (2023).

12. Halegua, T. et al. Delivery of Prime editing in human stem cells using pseudoviral NanoScribes particles. Nat Commun 16, 397 (2025).

13. Hamilton, J.R. et al. In vivo human T cell engineering with enveloped delivery vehicles. Nat Biotechnol 42, 1684–1692 (2024).

14. Hamilton, J.R. et al. Targeted delivery of CRISPR-Cas9 and transgenes enables complex immune cell engineering. Cell Rep 35, 109207 (2021).

15. Indikova, I. & Indik, S. Highly efficient ‘hit-and-run’ genome editing with unconcentrated lentivectors carrying Vpr.Prot.Cas9 protein produced from RRE-containing transcripts. Nucleic Acids Res 48, 8178-8187 (2020).

16. Ling, S. et al. Customizable virus-like particles deliver CRISPR-Cas9 ribonucleoprotein for effective ocular neovascular and Huntington’s disease gene therapy. Nat Nanotechnol 20, 543–553 (2025).

17. Lu, Z. et al. Lentiviral Capsid-Mediated Streptococcus pyogenes Cas9 Ribonucleoprotein Delivery for Efficient and Safe Multiplex Genome Editing. CRISPR J 4, 914–928 (2021).

18. Lyu, P., Javidi-Parsijani, P., Atala, A. & Lu, B. Delivering Cas9/sgRNA ribonucleoprotein (RNP) by lentiviral capsid-based bionanoparticles for efficient ‘hit-and-run’ genome editing. Nucleic Acids Res 47, e99 (2019).

19. Lyu, P., et al. Adenine Base Editor Ribonucleoproteins Delivered by Lentivirus-Like Particles Show High On-Target Base Editing and Undetectable RNA Off-Target Activities. CRISPR J 4, 69–81 (2021).

20. Lyu, P., Wang, L. & Lu, B. Virus-Like Particle Mediated CRISPR/Cas9 Delivery for Efficient and Safe Genome Editing. Life (Basel) 10 (2020).

21. Mangeot, P.E. et al. Genome editing in primary cells and in vivo using viral-derived Nanoblades loaded with Cas9-sgRNA ribonucleoproteins. Nat Commun 10, 45 (2019).

22. Raguram, A., An, M., Chen, P.Z. & Liu, D.R. Directed evolution of engineered virus-like particles with improved production and transduction efficiencies. Nat Biotechnol 43, 1635–1647 (2025).

23. Xu, D., et al. Programmable epigenome editing by transient delivery of CRISPR epigenome editor ribonucleoproteins. Nat Commun 16, 7948 (2025).

24. Barnes, C.R. et al. Genome-wide activation screens to increase adeno-associated virus production. Mol Ther Nucleic Acids 26, 94–103 (2021).

25. Iaffaldano, B.J., Marino, M.P. & Reiser, J. CRISPR library screening to develop HEK293-derived cell lines with improved lentiviral vector titers. Front Genome Ed 5, 1218328 (2023).

26. O’Driscoll, E.E., Arora, S., Lang, J.F., Davidson, B.L. & Shalem, O. CRISPR screen reveals modifiers of rAAV production including known rAAV infection genes playing an unexpected role in vector production. Mol Ther Methods Clin Dev 33, 101408 (2025).

27. Tridgett, M. et al. Lentiviral vector packaging and producer cell lines yield titers equivalent to the industry-standard four-plasmid process. Mol Ther Methods Clin Dev 32, 101315 (2024).

28. Han, J., Tam, K., Tam, C., Hollis, R.P. & Kohn, D.B. Improved lentiviral vector titers from a multi-gene knockout packaging line. Mol Ther Oncolytics 23, 582–592 (2021).

29. Xinyue, Z. et al. Engineering of HEK293T Cell Factory for Lentiviral Production by High-Throughput Selected Genes. CRISPR J 7, 272–282 (2024).

30. Lam, B. et al. Multi-species genome-wide CRISPR screens identify conserved suppressors of cold-induced cell death. eLife, 10.7554/eLife.102310.102311 (2024).

31. Replogle, J.M. et al. Combinatorial single-cell CRISPR screens by direct guide RNA capture and targeted sequencing. Nat Biotechnol 38, 954–961 (2020).

32. Seczynska, M., Bloor, S., Cuesta, S.M. & Lehner, P.J. Genome surveillance by HUSH-mediated silencing of intronless mobile elements. Nature 601, 440–445 (2022).

33. Seczynska, M. & Lehner, P.J. The sound of silence: mechanisms and implications of HUSH complex function. Trends Genet 39, 251–267 (2023).

34. DeKelver, R.C. et al. Functional genomics, proteomics, and regulatory DNA analysis in isogenic settings using zinc finger nuclease-driven transgenesis into a safe harbor locus in the human genome. Genome Res 20, 1133–1142 (2010).

35. Liao, J. et al. Therapeutic adenine base editing of human hematopoietic stem cells. Nat Commun 14, 207 (2023).

36. Johnson, S.S., Zhang, C., Fromm, J., Willis, I.M. & Johnson, D.L. Mammalian Maf1 is a negative regulator of transcription by all three nuclear RNA polymerases. Mol Cell 26, 367–379 (2007).

37. Vannini, A. et al. Molecular basis of RNA polymerase III transcription repression by Maf1. Cell 143, 59–70 (2010).

38. Vorlander, M.K. et al. Structural basis for RNA polymerase III transcription repression by Maf1. Nat Struct Mol Biol 27, 229–232 (2020).

39. Musunuru, K. et al. In vivo CRISPR base editing of PCSK9 durably lowers cholesterol in primates. Nature 593, 429–434 (2021).

40. Levy, J.M. et al. Cytosine and adenine base editing of the brain, liver, retina, heart and skeletal muscle of mice via adeno-associated viruses. Nat Biomed Eng 4, 97–110 (2020).

41. Bauler, M. et al. Production of Lentiviral Vectors Using Suspension Cells Grown in Serum-free Media. Mol Ther Methods Clin Dev 17, 58–68 (2020).

42. Richter, M.F., et al. Phage-assisted evolution of an adenine base editor with improved Cas domain compatibility and activity. Nat Biotechnol 38, 883–891 (2020).

43. Neugebauer, M.E., et al. Evolution of an adenine base editor into a small, efficient cytosine base editor with low off-target activity. Nat Biotechnol 41, 673–685 (2023).

44. Ngo, W. et al. Mechanism-guided engineering of a minimal biological particle for genome editing. Proc Natl Acad Sci U S A 122, e2413519121 (2025).

45. Liang, X., et al. Engineering of extracellular vesicles for efficient intracellular delivery of multimodal therapeutics including genome editors. Nat Commun 16, 4028 (2025).

46. Chen, Y.H. et al. Rapid Lentiviral Vector Producer Cell Line Generation Using a Single DNA Construct. Mol Ther Methods Clin Dev 19, 47–57 (2020).

47. Lee, Z., Lu, M., Irfanullah, E., Soukup, M. & Hu, W.S. Construction of an rAAV Producer Cell Line through Synthetic Biology. ACS Synth Biol 11, 3285–3295 (2022).

48. Powers, A.D., Drury, J.E., Hoehamer, C.F., Lockey, T.D. & Meagher, M.M. Lentiviral Vector Production from a Stable Packaging Cell Line Using a Packed Bed Bioreactor. Mol Ther Methods Clin Dev 19, 1–13 (2020).

49. Montagna, C. et al. VSV-G-Enveloped Vesicles for Traceless Delivery of CRISPR-Cas9. Mol Ther Nucleic Acids 12, 453–462 (2018).

50. Kunitake, K. et al. Barcoding of small extracellular vesicles with CRISPR-gRNA enables comprehensive, subpopulation-specific analysis of their biogenesis and release regulators. Nat Commun 15, 9777 (2024).

51. Lu, A. et al. Genome-wide interrogation of extracellular vesicle biology using barcoded miRNAs. Elife 7 (2018).

52. Strebinger, D. et al. Cell type-specific delivery by modular envelope design. Nat Commun 14, 5141 (2023).

53. Clement, K. et al. CRISPResso2 provides accurate and rapid genome editing sequence analysis. Nat Biotechnol 37, 224–226 (2019).

54. Shen, W., Le, S., Li, Y. & Hu, F. SeqKit: A Cross-Platform and Ultrafast Toolkit for FASTA/Q File Manipulation. PLoS One 11, e0163962 (2016).

55. Clement, K., Farouni, R., Bauer, D.E. & Pinello, L. AmpUMI: design and analysis of unique molecular identifiers for deep amplicon sequencing. Bioinformatics 34, i202–i210 (2018).

56. Langmead, B., Trapnell, C., Pop, M. & Salzberg, S.L. Ultrafast and memory-efficient alignment of short DNA sequences to the human genome. Genome Biol 10, R25 (2009).

57. Li, W. et al. MAGeCK enables robust identification of essential genes from genome-scale CRISPR/Cas9 knockout screens. Genome Biol 15, 554 (2014).

